# Deciphering the Signaling Network Landscape of Breast Cancer Improves Drug Sensitivity Prediction

**DOI:** 10.1101/2020.01.21.907691

**Authors:** Marco Tognetti, Attila Gabor, Mi Yang, Valentina Cappelletti, Jonas Windhager, Konstantina Charmpi, Natalie de Souza, Andreas Beyer, Paola Picotti, Julio Saez-Rodriguez, Bernd Bodenmiller

## Abstract

Although genetic and epigenetic abnormalities in breast cancer have been extensively studied, it remains difficult to identify those patients who will respond to particular therapies. This is due in part to our lack of understanding of how the variability of cellular signaling affects drug sensitivity. Here, we used mass cytometry to characterize the single-cell signaling landscapes of 62 breast cancer cell lines and five lines from healthy tissue. We quantified 34 markers in each cell line upon stimulation by the growth factor EGF in the presence or absence of five kinase inhibitors. These data – on more than 80 million single cells from 4,000 conditions – were used to fit mechanistic signaling network models that provide unprecedented insights into the biological principles of how cancer cells process information. Our dynamic single-cell-based models more accurately predicted drug sensitivity than static bulk measurements for drugs targeting the PI3K-MTOR signaling pathway. Finally, we identified genomic features associated with drug sensitivity by using signaling phenotypes as proxies, including a missense mutation in *DDIT3* predictive of PI3K-inhibition sensitivity. This provides proof of principle that single-cell measurements and modeling could inform matching of patients with appropriate treatments in the future.

**One-liner:** Single-cell proteomics coupled to perturbations improves accuracy of breast tumor drug sensitivity predictions and reveals mechanisms of sensitivity and resistance.

**HIGHLIGHTS:**

- Mass cytometry study of signaling responses of 62 breast cancer cell lines and five lines from healthy tissue to EGF stimulation with or without perturbation with five kinase inhibitors.
- Single-cell signaling features and mechanistic signaling network models predicted drug sensitivity.
- Mechanistic signaling network models deepen the understanding of drug resistance and sensitivity mechanisms.
- We identify drug sensitivity-predictive genomic features via proxy signaling phenotypes.

## INTRODUCTION

The aim of precision medicine is to use molecular markers of disease to enable tailored treatments. Currently, precision medicine is applied most often at the genetic level, in large part because genomic and transcriptomic measurements are scalable and cost effective. For example, tumors with the *BCR-ABL* fusion are usually successfully treated with imatinib mesylate (Gleevec), breast cancer with *HER2* overexpression is treated with trastuzumab (Herceptin), and melanomas that express BRAF^V600E^ are treated with vemurafenib (Zelboraf) (An et al., 2010; Garbe and Eigentler, 2018; Garrett and Arteaga, 2011). However, in treatment of breast cancer, patient-drug matching fails in a subset of patients, and, despite extensive characterization of genetic and epigenetic abnormalities in breast cancer, only a few targeted therapies are available (Coates et al., 2015; Network, 2012; Nik-Zainal et al., 2016; Pereira et al., 2016). Even a well-established biomarker like the amplification of *HER2* only partially predicts the tumor response: Only about half of all patients with *HER2*-amplified metastatic breast cancer respond to trastuzumab (Garrett and Arteaga, 2011).

Cancer cell lines have long been used as models for the human disease and to identify genomic features that correlate with and ultimately predict drug response (Barretina et al., 2012; Ghandi et al., 2019; Iorio et al., 2016; Neve et al., 2006). One aim of precision medicine is to identify and target the driver genomic alterations. Distinguishing passenger mutations from driver mutations remains challenging, however, some rare abnormalities are clearly oncogenic (Marcotte et al., 2016). Despite recent success in identifying driver alterations (Marcotte et al., 2016; Moghaddas Gholami et al., 2013), genomic information remains an incomplete predictor of drug sensitivity even in cell lines (Costello et al., 2014; Niepel et al., 2013). Genetic markers alone likely fail to predict drug response because genomic alterations have complex effects at the regulatory network and phenotypic level, and multiple drug resistance mechanisms at the level of signaling networks have been described (Lee et al., 2012; Yaffe, 2019). Phenotype-proximal readouts such as protein levels and post-translational modifications, which better reflect the status of the cell, are potentially better predictors of drug sensitivity than genomic sequence (Barrette et al., 2018; Beal et al., 2019; Fey et al., 2015; Fröhlich et al., 2018), especially when characterizing the response to a perturbation (Eduati et al., 2017; Hass et al., 2017; Meric-Bernstam et al., 2012; Niepel et al., 2013).

Many genetic and epigenetic alterations that drive cancer progression map to signaling pathways that control the key processes of growth, division, death, fate, metabolism, and motility (Forbes et al., 2011). Indeed, kinases and phosphatases involved in cellular signaling are the targets of some of the most effective anti-cancer therapeutics (e.g., HER2, EGFR, RAF) and some of the most promising future targets as well (e.g., PKC, p38, PI3K). However, the complex and redundant nature of the signaling network renders prediction of the effects of genomic alterations on the signaling state and drug sensitivity non-trivial. Furthermore, single-cell heterogeneity has been linked to fractional killing and drug resistance (Cooper and Bakal, 2017; Miura et al., 2018).

To develop a system to predict drug sensitivity, we used mass cytometry to map the single-cell signaling landscape of 62 breast cancer cell lines and five lines developed from healthy tissue. We quantified 34 markers over a 60-minute stimulation with the growth factor EGF in the presence or absence of five different kinase inhibitors. The generated breast cancer landscape revealed considerable heterogeneity in signaling responses at both the population and single-cell levels. Based on these multiparametric mass cytometric measurements, we built cell line-specific signaling network models. The cell line-specific models outperformed the state-of-the-art predictor based on transcriptomics data for PI3K-MTOR-targeting drugs. Finally, we identified genomic aberrations predictive of drug response that were not identified without the signaling characterization. Our analyses provide mechanistic insights into drug sensitivity and resistance mechanisms and suggest novel opportunities for patient stratification and combinatorial therapy.

## RESULTS

### THE PROTEOMES OF BREAST CANCER AND NORMAL BREAST CELL LINES

Since signaling networks are complex systems that can exhibit emergent properties, dynamic measurements under multiple conditions are required to model them effectively. As response to perturbation is known to be heterogeneous at the single-cell level and this heterogeneity is linked to drug resistance (Cooper and Bakal, 2017; Miura et al., 2018), we applied mass cytometry, using 35 antibodies (Sup. Table 1), to measure single-cell responses to EGF stimulation in the presence or absence of kinase inhibitors over a 10-points, 60-minute time course (Fig. 1, Sup. Table 2). The kinase inhibitors selected target key signaling nodes and are well-characterized and widely used: CI-1040 was the first MEK inhibitor to begin clinical development, pictilisib is a pan-PI3K inhibitor, rapamycin selectively inhibits mTOR, lapatinib inhibits both EGFR and HER2, and enzastaurin inhibits PKC (Allen et al., 2003; Folkes et al., 2008; Graff et al., 2005; Li et al., 2014; Xia et al., 2002) (Sup. Table 3). The resulting perturbation dataset includes quantitative information on 29 phosphorylation events covering the major signaling pathways, total protein abundance, DNA synthesis, and protein cleavage.

**Figure 1:**
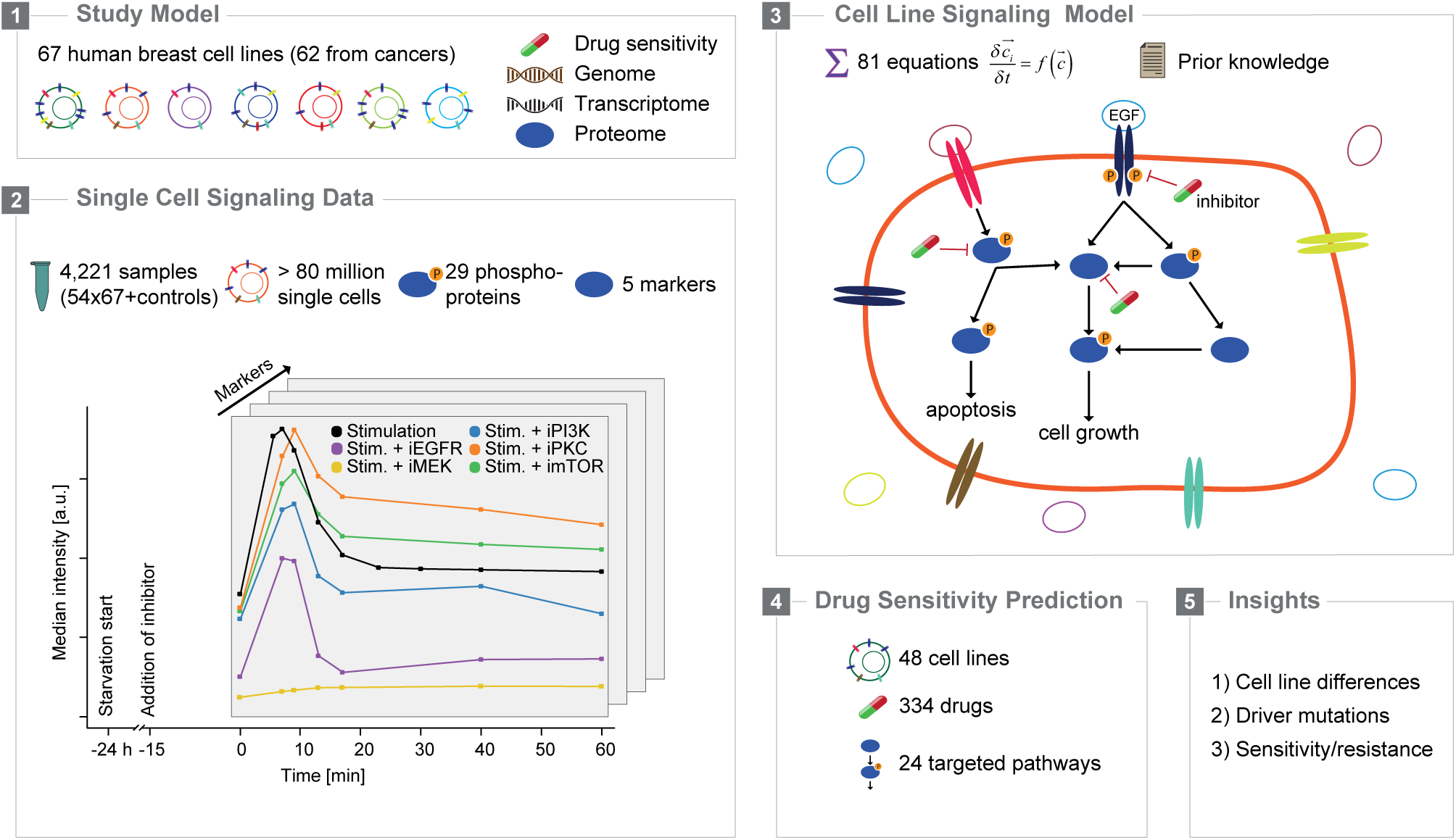
The Proteomes of Breast Cancer and Normal Breast Cell Lines. *Experimental and computational approach*.

We characterized the signaling landscapes of a panel of human breast cancer cell lines and cell lines from healthy breast tissue (Marcotte et al., 2016) (Sup. Table 4 and 5). The panel includes 62 cell lines generated from human breast tumors; 30 of these cell lines are basal-like and 32 are luminal-like, of which nine are known to overexpress HER2. These cell lines reflect the heterogeneity found in patient tumors, and both transcriptomic and genomic (single-nucleotide polymorphisms, SNPs, and copy number variations, CNVs) data are available for each cell line (Heiser et al., 2012; Marcotte et al., 2016; Neve et al., 2006). Importantly, for 48 of the cell lines in the panel, sensitivities (IC_50_ values) to 334 drugs have been measured (Picco et al., 2019; Yang et al., 2013). In total, we analyzed 4,000 samples and more than 80 million single cells (Fig. 1), making it the most comprehensive signaling response dataset to date.

Since there has been no systematic characterization of protein abundances for these cell lines, we first conducted a quantitative proteomic analysis of all cell lines in the panel using data-independent acquisition mass spectrometry. We quantitatively detected 9,031 proteins in cell lines grown without EGF stimulation. The proteomes of the five normal cell lines were very similar to each other (mean Pearson’s correlation coefficient for normal lines r = 0.94, across all lines r = 0.87) and clustered together (Sup. Fig. 1A). The good quantitative accuracy of the data is exemplified by the high correlation between levels of Ku70 and levels of Ku80 in all cell lines (Sup. Fig. 1B); the levels of these two proteins, which form a dimer involved in DNA double-strand break repair, are tightly controlled (Feng and Chen, 2012; Guo et al., 2019). The vast majority of detected proteins (7,328 proteins, 81%) were differentially abundant in at least one tumor cell line in comparison to the levels in the normal lines (Sup. Fig. 1C). On average, 2,600 proteins were differentially abundant when individual cancer cell lines were compared to the normal proteome; luminal cell lines had significantly more differentially expressed proteins than basal lines (Sup. Fig. 1D). In agreement with previous reports (Pozniak et al., 2016; Tyanova et al., 2016; Yanovich et al., 2018), the proteomes of luminal and basal cell lines mostly clustered together within each group and the separation between the two groups was mostly driven by the differential expression of proteins involved in metabolic processes and other known proteins such as FOXA1, Vimentin, CD44, HER2, MET, and EGFR (Sup. Fig. 1C, E, and F). The proteins that were differentially expressed between tumor and normal cell lines were enriched for breast-cancer-associated proteins and for GO-terms linked to cellular signaling (Sup. Fig. 1C and G), in agreement with prior knowledge that misregulated signaling plays an important role in cancer (Sanchez-Vega et al., 2018; Yaffe, 2019).

### THE SIGNALING LANDSCAPE OF BREAST CANCER CELL LINES

After analyzing static bulk proteomes of the cell lines, we exploited the dynamic single-cell data after EGF stimulation in order to examine the signaling responses by averaging phospho-protein levels across cells. Twenty-one of the measured markers significantly changed over time (ANOVA, adj. p-value ≤ 0.05). p-MEK^S221^, p-ERK, p-AKT^S473^, and p-S6 responded as expected (Fig. 2A) (Klinger et al., 2014; Pennock and Wang, 2003). A detailed examination of ERK-MAPK pathway markers revealed delayed peak times and signal amplification for proteins progressively more distal from the stimulus (Fig. 2B). Although abundances of most phosphorylated proteins increased upon stimulation, p-RB and p-4EBP1 levels decreased (Fig. 2A).

**Figure 2:**
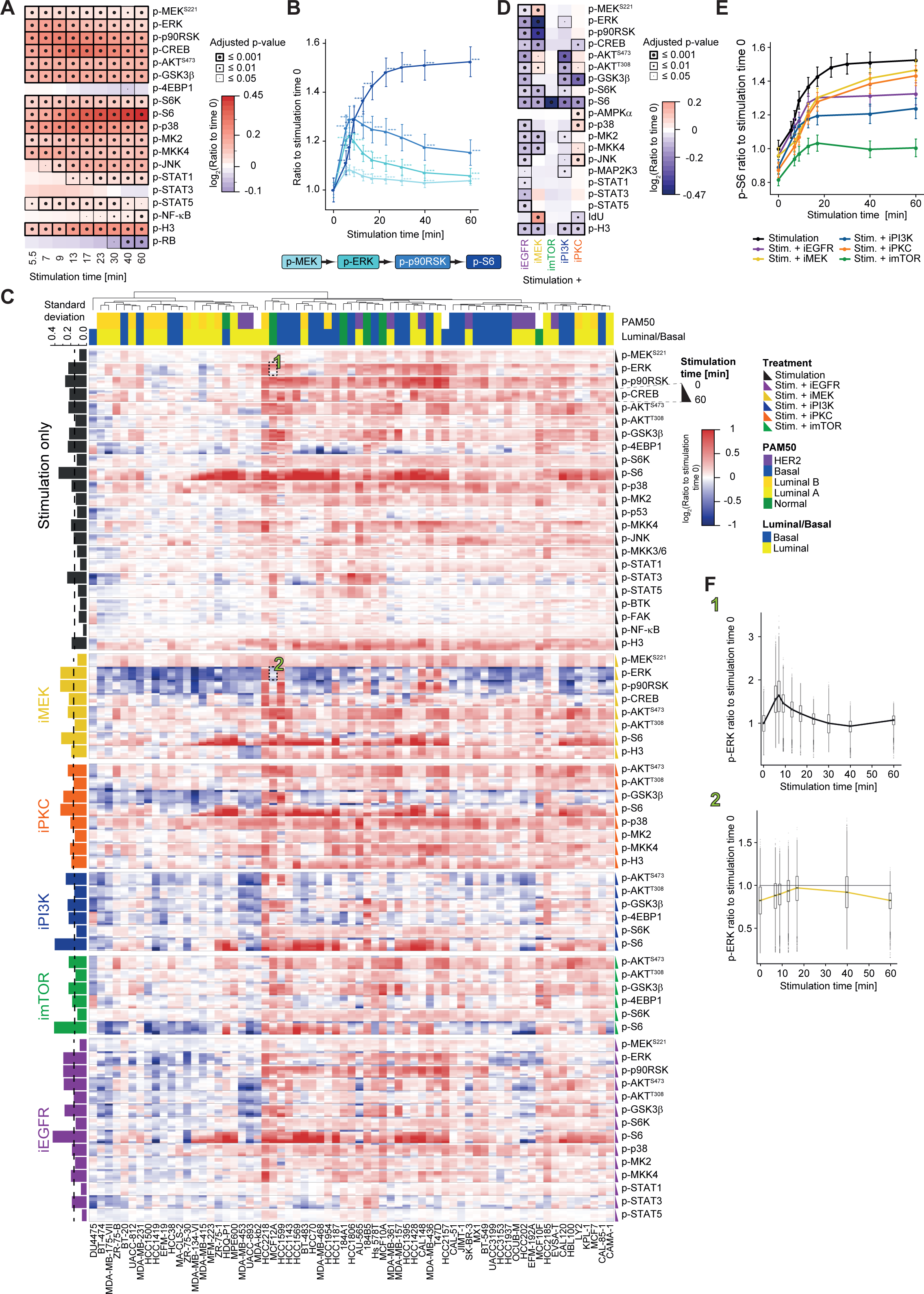
The Signaling Landscape of Breast Cancer Cell Lines. *(A) Median intensity ratios of markers to time point zero for markers with significant differences over time in response to stimulation with EGF when responses of the 67 cell lines are averaged. Adjusted p-values relative to time zero are represented by the dot size and the box thickness*. *(B) Ratio of signal at stimulation time vs. signal at time zero for indicated markers averaged over the 67 cells lines. The schematic depicts how signal is transmitted through the pathway. The error bars represent the standard error, and the asterisks the adjusted p-values (*p ≤ 0.05, **p ≤ 0.01, ***p ≤ 0.001)*. *(C) Ratios of marker abundance at a given time point compared to time zero, ordered by increasing stimulation time, marker, and treatment, in all 67 cell lines clustered based on their signaling signature. PAM50 tumor subtype classifications and luminal/basal classifications are overlaid. Data are from two independent experiments combined by linear interpolation. The bar graph to the left of heat map shows marker standard deviations across cell lines; the dotted line shows the treatment average. The single cell data underlying the regions numbered 1 and 2 are shown in panel F*. *(D) Median intensity ratios of markers significantly altered by the indicated kinase inhibitors compared to EGF stimulation alone. Adjusted p-values relative to time zero are represented by the dot size and the box thickness*. *(E) Ratios of phospho-S6 signal at the indicated EGF stimulation time vs. signal at time zero averaged over the 67 cells lines in the presence of the indicated kinase inhibitors. The error bars represent the standard error*. *(F) Values for 1) p-ERK signal in MCF12A cells upon EGF stimulation and 2) p-ERK signal in MCF12A cells upon EGF stimulation in the presence of the MEK inhibitor normalized to the average signal at time point zero and plotted against time. The medians correspond to those in the heat map shown in panel C and are indicated by thick lines. The 25% and 75% quantiles are indicated by the boxes. The whiskers extend between the median and ± (1.58 * inter-quantile range). Values beyond the whiskers are plotted individually*.

In individual cell lines, there were considerable differences in fold changes of all 34 measured markers upon EGF stimulation (Fig. 2C). Most cell lines responded strongly to EGF stimulation, but some did not respond at all and some even had lower levels of phosphorylation upon EGF stimulation (DU4475, BT-474, ZR-75-B, and MDA-MB-175-VII cells). These differences were not due to differences in initial levels of phosphorylation (Sup. Fig. 2A). Overall, p-NF-κB varied the least, and p-S6 and p-4EBP1 varied the most. Depending on the cell line, p-4EBP1 increased (HCC2218, MCF12A, HCC1599 cells) or decreased (MDA-MB-468, HCC1954, HCC1187 cells). In many cell lines there was no change in p-S6, but in 16 cell lines there was at least a 2-fold increase. Furthermore, phosphorylation of the STATs was highly cell line specific. Much of the inter-cell line heterogeneity was correlated. For example, p-ERK and p-p90RSK levels were correlated as were the two AKT phosphorylations, presumably due to common regulatory mechanisms. However, in certain cell lines this was not the case, hinting at differential regulatory mechanisms (Sup. Fig. 2B and C). Signaling dynamics varied between cell lines as well. In most lines, p-ERK peaked at 9 minutes and then decreased (Fig. 2B), but in some lines it had very different dynamics. For example, in T47D cells, levels plateaued at 9 minutes. Similar heterogeneity was observed for p-AKT^S473^, p-p90RSK, p-S6, and p-MKK4.

When the responses of all 67 cell lines were averaged, most of the measured markers changed significantly upon treatment with at least one of five kinase inhibitors when compared to EGF stimulation alone (20 markers, Fig. 2D). We observed an overall decrease in phosphorylation upon kinase inhibition, although certain markers increased, and all the inhibitors had the expected effects. For example, inhibition of EGFR resulted in reduced phosphorylation in the STAT, ERK-MAPK (p-MEK^S221^, p-ERK, p-p90RSK, p-CREB), and PI3K-AKT (both p-AKT sites, p-GSK3β, p-S6K, p-S6) pathways. Interestingly, both levels and dynamics of the known signal integrator S6 changed significantly upon inhibition of all five pathways (Fig. 2D and E). Analyses of average responses showed some intriguing behaviors. For instance, MEK inhibition induced an increase in levels of p-AKT but a decrease in p-S6K. Inhibition of PKC resulted in higher phosphorylation levels of several markers, including p-AMPKα, p-p38, and p-AKT^T308^, possibly due to the release of the PKC-mediated inhibition of the kinase GSK3β.

Notably, there was more heterogeneity between cell lines upon perturbation with kinase inhibitors than with EGF stimulation alone (Fig. 2C): The average standard deviation of the median fold change was significantly higher at 60 minutes upon treatment with all inhibitors than with EGF stimulation alone (ANOVA, adjusted p-value = 0.026). In some lines, the expected targets of inhibitors did not respond. Among the most interesting cases given the canonical signaling pathways were the cell lines in which ERK phosphorylation was observed despite the presence of the MEK inhibitor (e.g., T47D, HCC2218, HCC1599, CAL148 cells), GSK3β inhibition was observed although a PKC inhibitor was present (MCF12A, HCC2185, MCF10F cells), and p-AKT^S473^ phosphorylation was observed despite PIK3 inhibition (HCC2118, HCC1599 cells). In another example, inhibition of mTOR strongly inhibited phosphorylation of S6 in most cell lines at 60 minutes (mean decrease of 1.65 fold), but in MCF10F and HCC2185 cells (Fig. 2C and E). Overall, inhibition of EGFR resulted in the most unexpected behaviors in individual cell lines: We observed strong EGFR-dependent phosphorylation of S6 in MDA-kb2 cells and EGFR-independent phosphorylation of STAT3 and p90RSK in 184B5 and HCC202 cells, respectively. These phenotypes might be the result of either acquired resistance to the inhibitor or compensatory mechanisms.

When single cells from individual cell lines were evaluated, we generally observed homogenous responses to perturbations, and bimodal responses were rare. Cellular variability (quantified as the coefficient of variation of the different marker levels) decreased with EGF stimulation time but increased upon kinase inhibition (Sup. Fig. 2D and E). This phenomenon was apparent, for example, for p-ERK in MCF12A cells (Fig. 2F). There were differences among the cell lines, however. For instance, whereas p-S6 cellular variability typically decreased over time post EGF addition, this did not hold true for HCC1500 cells (Sup. Fig. 2F). In summary, some signaling patterns are clearly conserved across cell lines, but there were no two cell lines where the responses to inhibition were the same, revealing the width of the signaling landscape in breast cancer cell lines.

### CELL LINE-SPECIFIC SIGNALING NETWORK MODELS

Next, we used the generated data to train cell line-specific mechanistic signaling network models as a step toward understanding the signaling landscape of breast cancer cell lines. We began with the markers targeted by our antibody panel, expanded and connected the network using prior knowledge available in Omnipath (Türei et al., 2016) (Sup. Table 6, Sup. Fig. 3A), and built a dynamic mechanistic model using logic-based ordinary differential equations (Sup. Fig. 3B). For the cell line-specific signaling network models, we fit a node-specific speed factor (τ) that describes how rapidly the signal is relayed to that node via all upstream edges and an edge-specific transmission parameter (κ_A.B_) that non-linearly describes how much of the signal is relayed from node A to node B (Fig. 3A). Multiple steps are condensed into each of these parameters to increase scalability and to efficiently model multiple pathways together, which is required to use the complete set of markers. As a consequence, neither τ nor κ_A.B_ are directly interpretable in a biochemical sense; however, they provide measures of pathway activity. For example, a small node-specific speed factor τ cannot be interpreted as describing an enzyme with slow kinetics but would be expected for nodes with slow or minimal responses to a perturbation.

**Figure 3:**
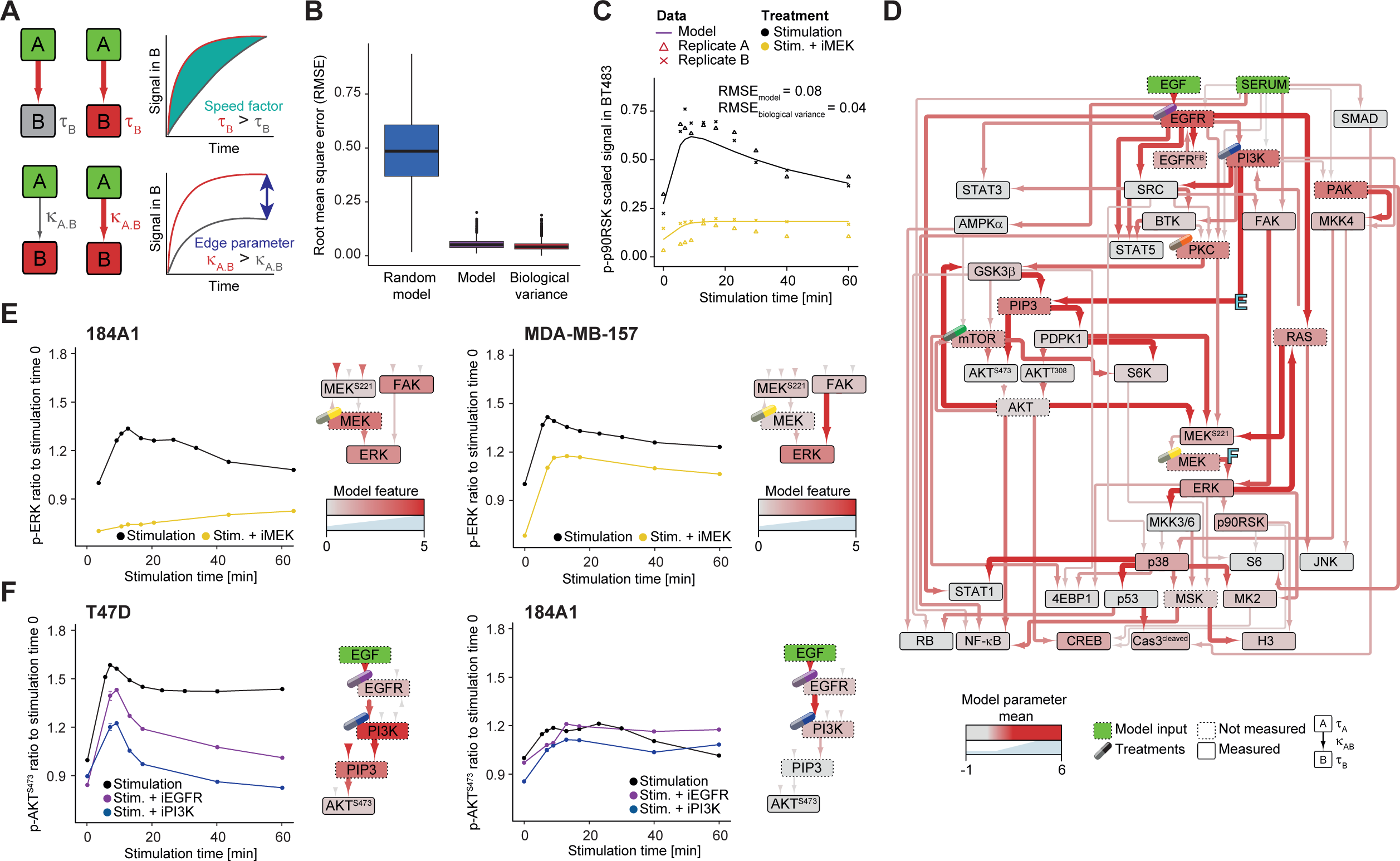
Cell-Line Specific Signaling Models. *(A) Illustrations of the effects of τ (speed parameter, top) and κ (edge parameter, bottom) on signal strength and dynamics in node B. The value of node A (input) changes over time from 0 to 1, and the signal of B is plotted as a function of time in the different modeled contexts depicted in the schematic*. *(B) Marker and cell line RMSE of a random model, the cell line-specific models, and the biological variance. The biological variance was computed as the average RMSEs between the medians and the two biological replicates for each marker and cell line. The thick lines indicate the median; the boxes and whiskers represent the 25% and 75% quantiles and the medians ± (1.58 * inter-quantile range), respectively. Data beyond the whiskers are plotted as dots*. *(C) Representative fit for the 85th percentile of the RMSE for the p-p90RSK signal upon stimulation with EGF without (black) and with MEK inhibition (yellow) in BT483 cells. The scaled signals for the biological replicates A and B are plotted as triangles and crosses, respectively. The fitted model is plotted as a continuous line*. *(D) The mean values for κ and τ of the mechanistic signaling network models for all 67 cell lines are represented as a signaling network. The color and thicknesses of edges indicate κ parameter values on a low-to-high scale (gray-red, thin-thick) and the node colors indicate τ parameter values on a low-to-high scale (gray-red). Modeled but not measured nodes are represented by dotted boxes, the model inputs are green, and intervention points are marked by an image of a drug capsule*. *(E) p-ERK signal over an EGF stimulation time course for 184A1 cells (left) and MDA-MB-157 cells (right) under the indicated conditions. The ratios of median signal to signal at time point zero are plotted. Error bars are standard errors of the median of single cells. The schematics in each plot show excerpts from the cell-line specific signaling model, represented as in panel D*. *(F) p-AKT^S473^ signal over an EGF stimulation time course for T47D cells (left) and 184A1 cells (right) under the indicated conditions. The ratios of median signal to that at time point zero are plotted. Error bars are the standard errors of the median of single cells. The schematics show relevant excerpts from the cell-line specific signaling model, depicted as in panel D*.

We tested the fits of models to the data using the root mean square error (RMSE); the lower the RMSE value the better the fit (Hengenius et al., 2014). The models fit the data very well with an average error of only 5%, which is in the same range as biological replicate average error of 4% (Fig. 3B). A few markers in some cell lines showed considerable error (0.3% of the marker and cell line combinations have a RMSE > 15%, and 5% with RMSE > 10%, e.g. 20% for p-S6K in AU565 cells and 17% for p-S6 in MPE600 cells), likely because the prior knowledge network is incomplete. Importantly, a single model for all cell lines performed poorly (data not shown), probably because it does not account for the observed heterogeneity between cell lines. The models captured both dynamics and inhibitor effects, as exemplified by the model for BT483 cells: The model describes accurately p-p90RSK time-dependent response to stimulation as well as its MEK-dependence (Fig. 3C). Clustering of the cell lines based on τ and κ partly recapitulated the major clusters obtained based on the response to stimulation (Fig. 2E, Sup. Fig. 3C). This suggests that our models – with only 107 model parameters – recapitulate the underlying 1,995 median points of information (markers x time points x treatments) in a condensed manner.

The average network across all cell lines, although not informative about cell-line heterogeneity, provides a compact view of how breast cancer cells process information (Fig. 3D). The most active pathways (as assessed by phosphorylation levels) have large signal transmission parameters. The highest κ was for the GSK3β·PIP3 edge (mean 3.14); this connection was one of the most consistently active across cell lines as it had the smallest coefficient of variation (CV = 67%, Sup. Fig. 3D). Other very active connections were PI3K·PIP3, EGF·EGFR, AKT·MEK^S221^, PIP3·AKT^S473^, p38·STAT1, ERK·MKK3, and ERK·MKK6. Furthermore, under the studied conditions, the activation of p38, S6, and CREB occurred mostly independently from their known activators MKK3, MKK6, and p90RSK (Fig. 3D), respectively (Remy et al., 2010; Roux et al., 2007; Xing et al., 1996).

The node-specific speed parameter τ is an indicator of reaction dynamics. Among the nodes with rapid dynamics in most cell lines were EGFR, PI3K, PIP3, PAK, and PKC (Fig. 3D). In contrast, SMAD2, SMAD3, AMPKα, the STATs, SRC, MKK3, MKK6, and p53 nodes generally had slow dynamics (Fig. 3D). The parameter τ is highly cell line dependent with a mean CV of 164%. The most conserved τ was that for the GSK3β to PIP3 edge with a CV of 67% (Sup. Fig. 3D). The parameters weakly correlated across cell lines (Pearson’s correlation, *r* = 0.17, Sup. Fig. 3E), reflecting the heterogeneous signaling landscape. Models of some cell lines were more correlated, indicative of quite similar dynamics (*r* = 0.61 for HCC1428 and MDA-MB-362 cells), whereas others were very different (*r* = −0.15 for HCC1599 and EFM-192A cells, Sup. Fig. 3E). Transmission parameters, κ, were significantly less variable than τ across cell lines (16 of 88 had a CV under 100%, Sup. Fig. 3D and F).

Since MEK is a clinical target of many drugs and MEK-independent ERK activation is a known resistance mechanism (Grimaldi et al., 2017; Kim and Giaccone, 2018; Simard et al., 2015), we examined MEK pathway activity. In most cell lines this pathway was active; however, there were some striking differences. For example, in 184A1 cells, ERK activation was mostly MEK dependent, whereas in MDA-MB-157 cells it was mostly FAK dependent (Fig. 3E). Our data are in line with previous reports that show that MDA-MB-157 cells are relatively resistant to BRAF-targeted drugs (PLX-4720 and dabrafenib) in comparison to the other cell lines (Picco et al., 2019; Yang et al., 2013). Cell line-specific differences were also observed in the generally active PI3K pathway: In T47D cells, the phosphorylation of AKT^S473^ depended strongly on PI3K, whereas in 184A1 cells the dependency was less pronounced (Fig. 3F). This is consistent with the fact that T47D cells are sensitive to PI3K inhibition (Picco et al., 2019; Yang et al., 2013), 184A1 was not included in this study. These cell line-specific differences could be indicative of opportunities for intervention or patient stratification.

### PREDICTION OF DRUG SENSITIVITY USING DYNAMIC PREDICTORS

We used the Genomics of Drug Sensitivity in Cancer (GDSC) dataset which includes IC_50_ values for 334 drugs in 48 of the cell lines in our panel (Picco et al., 2019; Yang et al., 2013), to assess whether our dynamic models accurately predicted drug sensitivity. We used machine learning to predict the IC_50_ values using either static or dynamic predictors (Fig. 4A). The static predictors, describing the steady state and acquired in absence of perturbation, included protein abundance measurements (*log_2_*fold change to the normal), RNA-seq data, and the non-linear logic-based model described in the previous section, parametrized by the τ and κ model parameters. Since the τ and κ parameters of the cell line-specific logic signaling network models are time- and treatment-independent, these parameters do not provide direct information on which pathways are active in specific conditions or on the covariance over time and are thus considered as static. The dynamic predictors, describing the perturbed state include 46 interdependent matrices, one for each individual combination of treatment and time point (Sup. Table 2), and include the median marker expression, the variability of marker expression at the single-cell level, and the edge flux. The edge fluxes are computed from the model parameters (τ and κ) and the median marker expression for a specific condition (treatment and time point) and represent the activity transferred between each node pairs similarly to metabolic fluxes.

**Figure 4:**
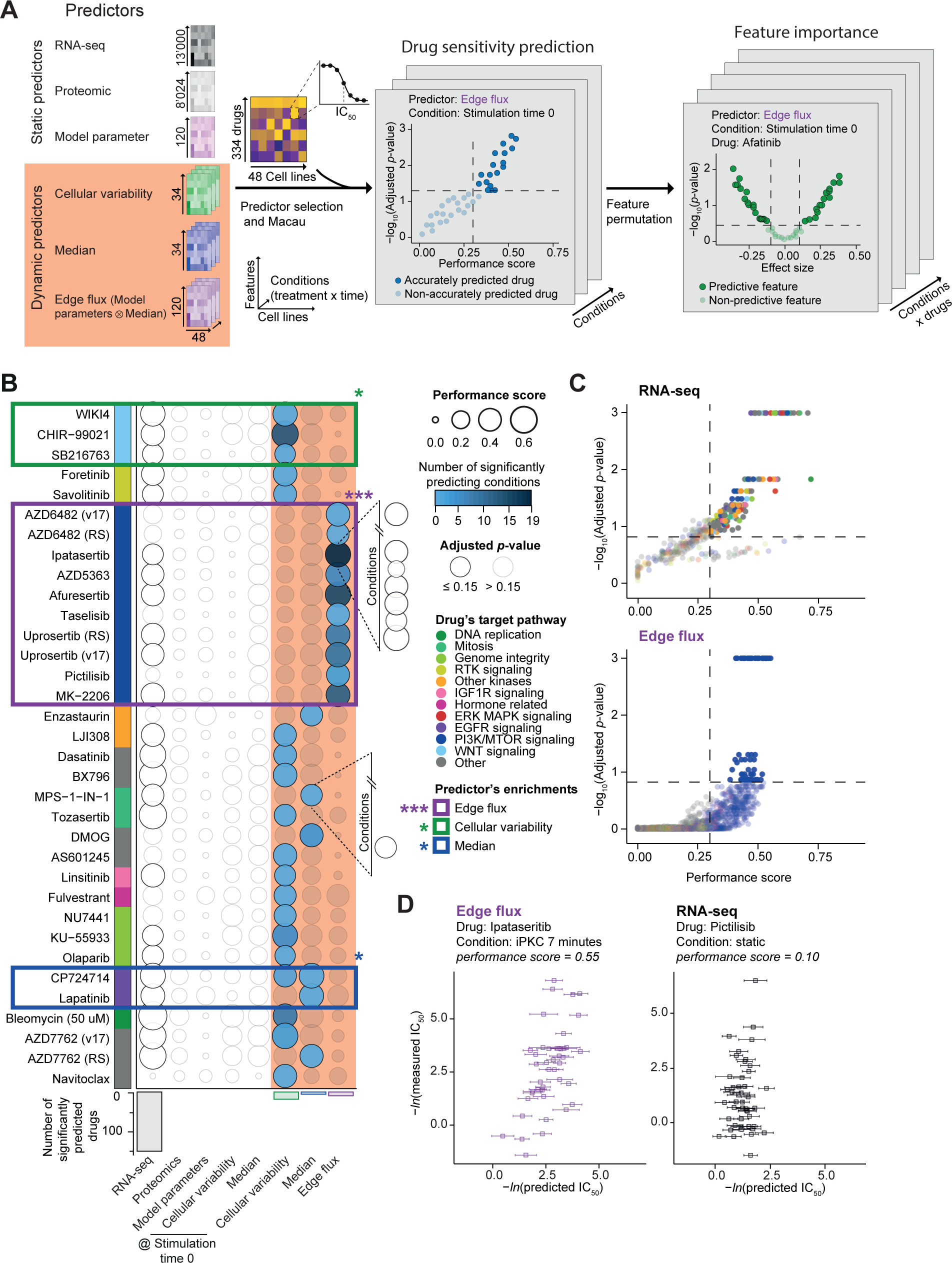
Prediction of Drug Sensitivity Using Dynamic Predictors. *(A) Computational approach to predict drug sensitivity and identify predictive features*. *(B) Upper: Sensitivities that are predicted with significant accuracy by at least one dynamic predictor are shown (FDR 15% and performance score > 0.3, multiple hypothesis correction for the predicted drug measurements, n = 409) in rows versus the predictors in columns. Cellular variability and median were used as both static (stimulation time zero) and dynamic predictors, shown separately. The bubble color indicates the number of times the drug sensitivity was predicted with significant accuracy (for 46 combinations of treatment and time). The bubble size is proportional to the performance score of the best predictor. If the bubble circumference is light gray, sensitivity was not accurately predicted. Drugs are arranged by their target pathways (key to the far right); significantly predicted pathways are marked with a colored box and p-values are shown at top right (*p ≤ 0.05, **p ≤ 0.01, ***p ≤ 0.001; Fisher’s exact test). Parentheses following the drug names give the version of the GDSC screen, if ambiguous. Lower: Number of accurately predicted drugs per predictor is reported as a bar plot*. *(C) Performance score plotted against the significance for predictions using the RNA-seq (top) and edge flux (bottom) predictors. Color code indicates putative target pathways. Thresholds for significance are indicated by dashed lines (FDR 15% and performance score > 0.3)*. *(D) Plots of measured and predicted IC_50_ values for the cell lines for which data are available from the GDSC dataset. Left: Ipataseritib sensitivity was best predicted by the edge flux at 7 minutes stimulation in presence of the PKC inhibitor. Right: Pictilisib sensitivity was not significantly predicted with RNA-seq data. The error bars represent the standard deviation from the 5-fold cross-validation*.

To predict drug sensitivity, we employed the Macau algorithm. Macau is a machine-learning approach based on scalable Bayesian multi-relational factorization with side information using Markov chain Monte Carlo (Simm et al., 2017; Yang et al., 2018) (Fig. 4A). We defined as a performance score the Pearson’s correlation between predicted and measured IC_50_ (in a cross-validation scheme, where predicted cell-lines are not used for training) and identified significantly predicted drug sensitivity by requiring an FDR of less than 15% and a performance score greater than 0.3. Based on these criteria, RNA levels (Marcotte et al., 2016) were predictive for the cell line sensitivity to 149 drugs (45%). Compared to RNA levels, variability of marker expression at the single-cell level, the edge flux, and median marker expression performed worse as predictors across drugs with various mechanisms (19, 9, and 6 drug sensitivities accurately predicted, respectively) (Fig. 4B). Use of protein abundance data, model parameters (τ, κ), or an RNA-seq dataset reduced to 34 dimensions with sparse principal component analysis (Erichson et al., 2018) predicted no sensitivities accurately (Fig. 4B, Sup. Fig. 4A and data not shown). As expected, larger IC_50_ ranges and less missing data yielded more significant predictions (Sup. Fig. 4B).

Whereas RNA-seq data accurately predicted sensitivities of more cell lines to drugs than other static inputs, there was no enrichment for drugs targeting common pathways (Fig. 4C). In contrast, the drug sensitivities accurately predicted by the dynamic predictors (edge flux, cellular variability, and median marker expression) were significantly enriched for drugs targeting parts of the modeled network (Fisher’s exact test, Fig. 4B). The edge flux most accurately predicted sensitivities of cell lines to drugs targeting the PI3K-MTOR signaling pathway (Fig. 4C). For instance, sensitivities to ipataserib and pictilisib were better predicted by the edge flux than by the RNA-seq-based model, and only the edge flux predicted sensitivities to pictilisib with significant accuracy (Fig. 4D and Sup. Fig. 4C). In contrast, cellular variability and median marker expression were the most accurate predictors of sensitivities to drugs targeting the WNT and EGFR signaling pathways, respectively (Fig. 4B).

For the 46 interdependent edge flux matrices, we plotted how many combinations of inhibitor treatment and time (i.e., conditions) were predictive of drug sensitivity (Fig. 4B). In general, only a few conditions were predictive of sensitivity to a given drug (median of 1 condition per drug, Sup. Fig. 4A). The edge flux was the most consistent predictor across conditions (median of 7 conditions per drug, e.g., the AKT-targeting drugs ipatasertib and afuresertib). The median marker expression and cellular variability were also predictive of sensitivity to some drugs across conditions; however, these predictors were not accurate when only the stimulation condition was considered, demonstrating the importance of perturbation experiments (Sup. Fig. 4A).

For different drugs, the predictors that were accurate differed. Sensitivities of just two drugs (CP724714 and AZD7762) of the 33 significantly predicted by the dynamic predictors were accurately predicted by two predictors (cellular variability and median marker expression), showing that the predictors provide orthogonal information (Fig. 4B). For ten drugs (AS601245, AZD6482, DMOG, enzastaurin, fulvestrant, navitoclax, NU7441, pictilisib, and taselisib), sensitivities were accurately predicted only by the dynamic predictors. Thus, the characterization of the signaling landscape improved drug sensitivity prediction of selected kinase inhibitors.

### FEATURES AFFECTING DRUG SENSITIVITY PREDICTIONS

We next identified the features, such as the median expression of a phosphorylated protein, that are important for the drug-sensitivity prediction (Fig. 4A, right side). We extracted feature importance directly from the drug sensitivity models using a procedure similar to that used to retrieve loadings for linear models. We calculated an effect size per drug for each feature and computed the significance of each effect size (Fig. 4A). The effect size is a measure of the contribution of a particular feature to the accurate drug sensitivity prediction. We defined features with positive effects as those higher in resistant cell lines and not in sensitive ones; features with negative effects were those higher in sensitive but not resistant cell lines.

Features that significantly contributed to overall accuracy of prediction of sensitivity and resistance (effect size > 0.01 or < −0.01 and 5% FDR) were identified by averaging across all models for all drugs (Fig. 5A and Sup. Fig. 5A). The edge flux has more predictive features than other predictors (11 of the 15 most predictive features, Fig. 5B). For example, three of the edges connecting PKC to the signaling network in the contextual model are among the 15 most predictive features, with the edge from BTK to PKC being the most predictive feature. The most predictive feature is GAPDH. Whereas GAPDH median levels were predictive of sensitivity, its cellular variability was predictive of resistance. The opposing effects of median levels and cellular variability were also observed for other proteins such as Ki-67, p-MKK4, and p-MEK^S221^ (Sup. Fig. 5A and B).

**Figure 5:**
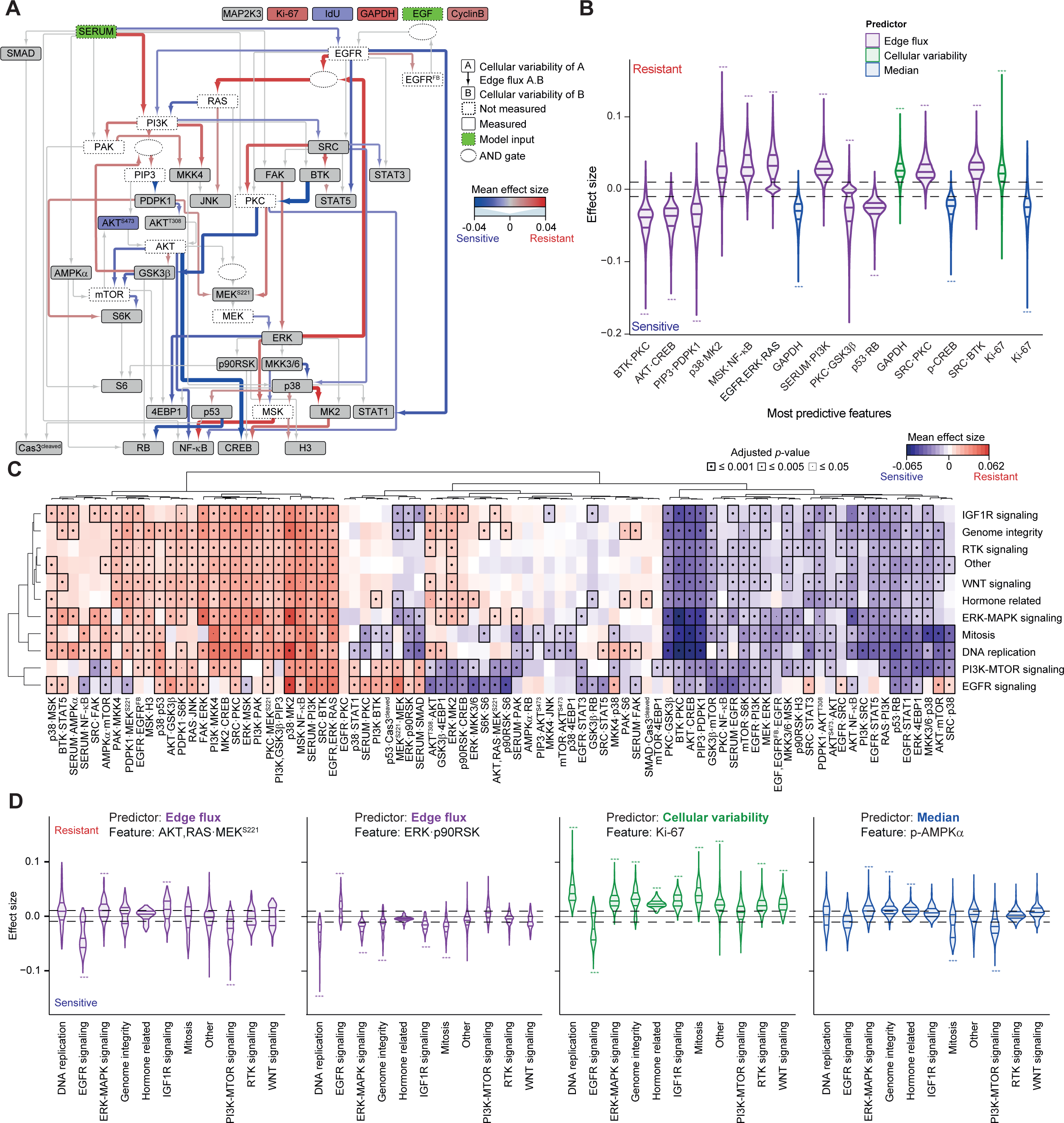
Features that Influence Drug Sensitivity Predictions. *(A) Significant effect size features of the cellular variability (nodes) and edge flux (edges) predictors are represented on the signaling network. The colors represent the mean effect sizes over all conditions and drugs. The edge thicknesses are proportional to the absolute values of the mean effect sizes (FDR 1%)*. *(B) Effect size distributions for the most predictive 16 features across the three dynamic predictors. The horizontal lines represent the medians and the 25% and 75% quantiles. The color of the distribution indicates the predictor. The effect size threshold (0.01) is indicated by the dashed line. *p ≤ 0.05, **p ≤ 0.01, ***p ≤ 0.001*. *(C) Mean pathway-specific effect size features of the edge flux with drugs binned according to the target pathway in rows and features shown in columns (FDR 5%). Both the features and target pathways were hierarchically clustered. Mean effect sizes are indicated on a low-to-high color scale. For each class the lowest adjusted p-value of all side-by-side comparisons is indicated by the dot size and the box thickness. The group “Other” contains all the drugs not falling into another group*. *(D) Effect size distributions for four selected features showing pathway-specific effects. The selected predictors and features are indicated in each case. The significant threshold of 0.01 minimum effect size is plotted as a dashed line. The horizontal lines indicate the median and the 25% and 75% quantiles. AKT,RAS·MEK^S221^ represents the parameter controlling MEK activation and it integrates both positive RAS and negative AKT influence. *p ≤ 0.05, **p ≤ 0.01, ***p ≤ 0.001*.

Next, we repeated the same analysis but instead of averaging across all drugs we averaged across groups of drugs targeting selected biological pathways (Fig. 5C, Sup. Fig. 5C and D). This approach minimized contributions of off-target effects and provided insights into drug-class specific effects. While most features with a significant predictive effect were correlated with sensitivity (e.g., BTK·PKC) or resistance (e.g., p38·MK2) across all drug classes (Fig. 5A and Sup. Fig. 5A), we observed drug class-specific patterns in which certain features are predictive only for drugs targeting a certain pathway (Fig. 5C, Sup. Fig. 5C and D). We also identified features that are sometimes predictive of sensitivity and sometimes of resistance, depending on the targeted pathway. For example, the AKT,RAS·MEK^S221^ edge flux (the parameter controlling MEK activation, which integrates both the positive influence of RAS and the negative influence of AKT) correlated with sensitivity to drugs targeting EGFR and PI3K-MTOR signaling and resistance to drugs targeting ERK-MAPK and IGF1R signaling (Fig. 5C and D). Although not predictive for overall drug sensitivity or resistance, p90RSK correlated with sensitivity to drugs targeting several specific pathways (Fig. 5A and C, Sup. Fig. 5A, C, and D): Its activation through ERK (ERK·p90RSK) was predictive of sensitivity to PI3K-MTOR pathway-targeted drugs, p90RSK·S6 of sensitivity to drugs targeting EGFR signaling, and its cellular variability of resistance to ERK-MAPK signaling drugs (Fig. 5D and Sup. Fig. 5E). In other examples, the cellular variability of p-STAT1 was particularly predictive for a subset of drugs that target PI3K-MTOR, and the median level of p-AMPKα was predictive of sensitivity to PI3K-MTOR inhibition (Sup. Fig. 5C and D). An interesting instance is the extent to which the cellular variability of the proliferation marker Ki-67 predicted resistance across drug classes (Fig. 5D and Sup. Fig. 5E). Large Ki-67 variability was predictive of sensitivity to EGFR signaling drugs. Importantly, cellular variability of Ki-67 was most predictive for drugs targeting DNA replication. This supports the “proliferation rate paradox”: that is, the finding that many chemosensitive human cancers have low proliferation rates (Mitchison, 2012). Interesting identified trends comprise the dependence on the source of activation (e.g., ERK, PI3K, and PKC, Fig. 5A) and the opposite effects on closely related pathways of some features (e.g. ERK·p90RSK opposite effects on EGFR and ERK-MAPK signaling, Fig. 5D). In sum, these findings confirm the utility of integration of dynamic perturbation data by mathematical modeling and show how identical signaling features can be predictive of sensitivity and resistance depending on the context.

### RESISTANCE AND SENSITIVITY TO PI3K AND EGFR INHIBITION

Finally, we used the dynamic predictors to probe mechanisms of drug resistance and sensitivity. We focused on inhibition of two clinically relevant targets, EGFR and PI3K, considered essential to basal and luminal breast cancers, respectively (Marcotte et al., 2016). The median marker expression after treatment with the PI3K inhibitor for 60 minutes predicted the sensitivities of cell lines to the FDA-approved lapatinib, an inhibitor of EGFR and HER2 used in combination therapy for HER2-positive breast cancer (Giampaglia et al., 2010), with significant accuracy (Fig. 4B). We identified the features significantly predictive of lapatinib resistance or sensitivity for all dynamic predictors, since they all performed quite well in predicting lapatinib sensitivity (Fig. 4B and Sup. Fig. 6A). Consistent with previous reports (Campbell et al., 2004; Zhang et al., 2011), alternative activation of PI3K (SERUM·PI3K, in contrast to EGF-dependent PI3K activation via EGFR·PI3K and RAS·PI3K) and SRC correlated with resistance, and strong EGFR/HER2 activity (SERUM·EGFR, modeling the EGF-independent EGFR activation) correlated with sensitivity (Fig. 6A). Strikingly, the median expression levels of one of the two phosphorylated forms of AKT correlated with resistance and the other with sensitivity. The variabilities of p-STAT5 and p-STAT3 were more predictive of sensitivity than were the median levels of expression (Fig. 6A and Sup. Fig. 6A); median expression of the phosphorylated forms of STAT5 and STAT3 were previously reported to be predictive of lapatinib, canertinib, and afatinib sensitivity (Gschwantler-Kaulich et al., 2016). Interestingly, p-MK2 median expression correlated with sensitivity, and the p38·MK2 and ERK·MK2 edge fluxes strongly correlated with resistance and sensitivity, respectively. These results are indicative of the importance of MK2 in the response of cells to lapatinib.

**Figure 6:**
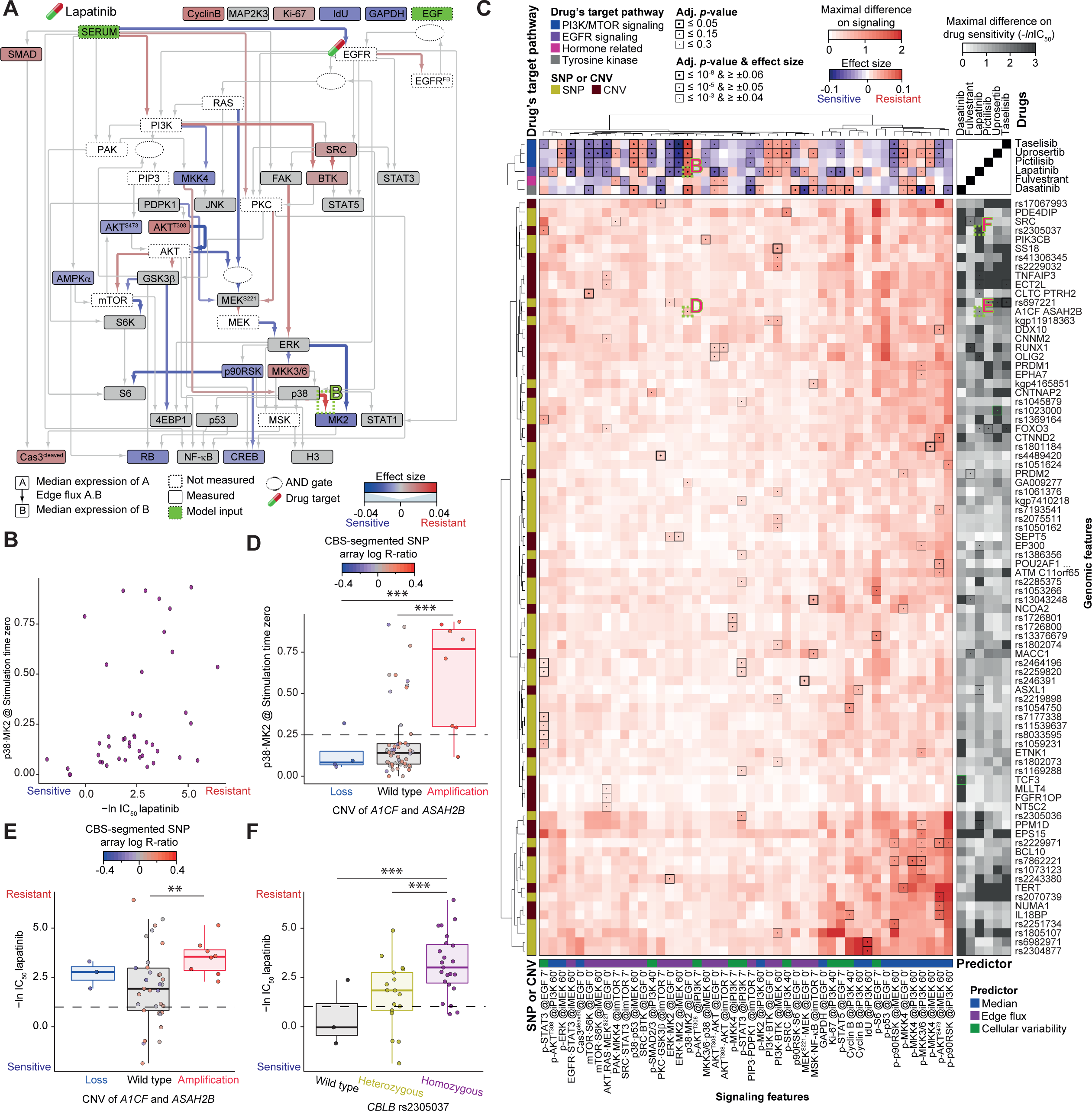
Resistance and Sensitivity to PI3K and EGFR Inhibition. *(A) Significant effect-size features of the median (nodes) and edge flux (edges) predictors represented on the signaling network for lapatinib (FDR 15% and effect size > 0.01 or < −0.01). The effect sizes are indicated using color (nodes and edges) and the edge thicknesses are proportional to the absolute values of the effect sizes. An image of a drug capsule highlights the putative drug target. In each case the conditions with the highest performance scores are plotted: PI3K inhibitor 60 minutes and starvation for medians and edge fluxes, respectively. The data in region labeled B is shown in that panel*. *(B) Values for the edge flux p38·MK2 (at stimulation time zero) plotted against lapatinib IC_50_ for different cell lines (n = 42). Each data point represents a cell line*. *(C) Upper: Heat map of the effect size of the selected signaling features (columns) and the six selected drugs (rows) for 48 cell lines. The drug’s target pathway and predictors are overlaid. The effect sizes are indicated on a low-to-high color scale. The adjusted p-value and effect size thresholds are indicated by the dot size and the box thickness. Lower left: Heat map of the maximal difference in selected signaling features (columns) among genomic statuses (rows) in 64 and 61 cell lines for filtered CNV and SNP, respectively. Both the signaling features and the genomic features were hierarchically clustered. SNP or CNV status is overlaid. Only genomic features with at least one significant link resulting from the QTL analyses are reported, and the adjusted p-values are indicated by the dot size and the box thickness. Lower right: Heat map of maximal differences in IC_50_ values (–ln) for dasatinib, fulvestrant, lapatinib, pictilisib, taselisib, and uprosertib (columns) among genomic statuses (rows) in 42 and 40 cell lines for CNVs and SNPs, respectively. The adjusted p-value of the QTL analysis using the IC_50_ values directly and using an ANOVA on the genomic features identified through the signaling are indicated by the dot size and the box thickness in green and black, respectively. In both the lower left and right heat maps, the maximal differences are indicated on a low-to-high color scale capped at two and three, respectively. The data in regions labeled B, D, E, and F are shown in those panels*. *(D) Values of the edge flux p38·MK2 (at stimulation time zero) plotted against CNV status of A1CF/ASAH2B (n = 64 cell lines). Thick lines indicate medians, the dashed line indicates the arbitrary threshold for high activity, boxes indicate the 25% and 75% quantiles, and whiskers extend between the median and ± (1.58 * inter-quantile range). Each data point represents a cell line, and the color intensity indicates the amplification status on a low-to-high scale (blue-gray-red, ANOVA followed by Tukey honest significant differences computation: *p ≤ 0.3, **p ≤ 0.15, ***p ≤ 0.05)*. *(E) IC_50_ values (–ln) for lapatinib plotted against CNV status of A1CF and ASAH2B (n = 42 cell lines). Plots and statistical analysis are as in panel E, except that the dashed line indicates the arbitrary threshold for sensitivity (1)*. *(F) IC_50_ values (–ln) for lapatinib plotted against SNP status of rs2305037 (CBLB gene, n = 40 cell lines). Plots and statistical analysis are as in panel E, except that the dashed line indicates the arbitrary threshold for sensitivity (1) and that colors indicate the SNP status*.

The edge flux was the only predictor that accurately identified cell lines sensitive to pictilisib, a pan-PI3K inhibitor (Fig. 4B, D and Sup. Fig. 4B). S6K activation by mTOR (mTOR·S6K), activation of STAT1 and STAT3 (EGFR·STAT3, SRC·STAT3, and EGFR·STAT1), and phosphorylation levels of S6K, STAT1, and STAT3 were sensitivity predictors, whereas induction of the oncogene-induced senescence pathway (p38·p53) was predictive of resistance (Sup. Fig. 6B). ERK pathway activation and cellular variability of the phosphorylated STATs were predictive of sensitivity to pictilisib (Sup. Fig. 6B), PI3K-MTOR inhibitors (Fig. 5C, Sup. Fig. 5C and D), and the PI3K-inhibitor taselisib (Sup. Fig. 6C). Pictilisib-specific effects were also observed: cellular variabilities of p-p53 were predictive of sensitivity to pictilisib but not to taselisib or the PI3K-MTOR drug group and EGFR·PKC was only prominently predictive of resistance to pictilisib (Fig. 5C, Sup. Fig. 6B and C). Off-target effects or differences in targets or mechanism could explain such drug-specific characteristics, and these must be considered when designing clinical trials.

Using dynamic predictors, we were also able to predict sensitivities of drugs that do not directly target a protein present in our network, likely because these drugs indirectly affect the network. For example, cellular variability was the only predictor of response to fulvestrant (Fig. 4B), a selective estrogen receptor degrader used to treat hormone receptor-positive breast cancer (Nathan and Schmid, 2017). Estrogen signaling interplays with both the ERK-MAPK and the PI3K-MTOR signaling pathways (Tanos et al., 2012), which were monitored by the antibodies in our mass cytometry panel. An analysis of feature importance revealed that EGFR, SRC, and ERK play defining roles in the response to this drug (Sup. Fig. 6D). Hence, our predictors can be used even when the direct targets are not measured, as long as they are regulated through the signaling pathways that we model.

The identification of genomic aberrations correlating with drug response is an essential step toward efficacious personalized medicine; genomic aberrations could serve as biomarkers and help revealing the mechanisms of drug resistance. However, their identification is challenging, and the effect sizes often limited (Garnett et al., 2012; Menden et al., 2018). Indeed, a direct quantitative trait locus (QTL) analysis (Clément-Ziza et al., 2014) of sensitivity to dasatinib, fulvestrant, lapatinib, pictilisib, taselisib, and uprosertib identified only one copy number variant (CNV) and one single nucleotide polymorphism (SNP) associated with sensitivity: *TCF3* for dasatinib and *FNBP1* (rs1023000) for uprosertib, respectively (Fig. 6C, green highlights on the right heat map). This was in spite of the restriction of the search space to the COSMIC list of oncogenes (736 SNPs and 518 CNVs in 706 oncogenes) (Sondka et al., 2018) and a 30% FDR threshold(Eduati et al., 2017). We wondered if by taking advantage of the signaling characterization we could improve the QTL performance; therefore, we ran QTL analyses on 46 signaling features predictive of sensitivity to dasatinib, fulvestrant, lapatinib, pictilisib, taselisib, and uprosertib (Sup. Table 7, Fig. 6C). The analysis revealed associations between signaling features and 55 SNPs and 38 CNVs (Fig. 6C). For instance, *A1CF*/*ASAH2B* CNV status was predictive of the lapatinib-predictive p38·MK2 edge flux (Fig. 6C and D) and the rs2305037 SNP (*CBLB* gene) of p-STAT3 cellular variability (Sup. Fig. 6E). Interestingly, although we did not identify any genomic aberrations linked to lapatinib sensitivity directly, the signaling characterization enabled to discover that cell lines with amplification in *A1CF*/*ASAH2B* or those homozygous for rs2305037 were generally more resistant to lapatinib (Fig. 6E and F). Furthermore, we identified ten genomic aberrations linked to the median expression levels of p-MKK3 and p-MKK6, which were, in turn, predictive of lapatinib resistance (Fig. 6A). Two of these genomic aberrations (*EP330* and *PPM1D*) showed potential in differentiating lapatinib response (Sup. Fig. 6F). In addition, CNVs of *PRDM2* and *MACC1* were linked to fulvestrant sensitivity through their link to p-MKK4 median expression and the MSK·NF-κB edge flux, respectively (Sup. Fig. 6G). Additionally, the contextualized feature ERK·MK2 was associated with the SNP rs697221, causing a missense change in the sequence of the gene *DDIT3*. ERK·MK2 was predictive of sensitivity to all three PI3K targeted inhibitors studied (pictilisib, uproseritib, and taselisib). Overall through the signaling characterization, we identified 21 genomic aberrations predictive of drug sensitivity (none were identified for dasatinib, Fig. 6C, black highlights on the right heat map). Thus, performing the QTL on the signaling phenotype can identify SNPs and CNVs predictive of drug sensitivity in cases where the correlation to drug sensitivity was too weak to be identified directly.

## DISCUSSION

Here, we report the largest single-cell signaling dataset collected to date: We used mass cytometry to interrogate signaling in a panel of 62 breast cancer cell lines and five cell lines generated from normal tissue with or without stimulation with EGF and in the presence or absence of five kinase inhibitors. We also characterized the proteomes of these cell lines in unstimulated conditions. Although clustering of the proteomic data from unstimulated cells resulted in accurate separation of the cancerous lines into luminal and basal phenotypes, in both the signaling and model parameter clustering there was not a clear basal/luminal separation. We did, however, observe a luminal-enriched cluster of cell lines that were less responsive to kinase inhibitor treatment than the average cell line. Cell line-specific network models showed that PI3K pathway activation and rapid EGFR negative feedback characterize these non-responsive cell lines. The proteomic profiling data also showed that phosphorylation and collagen-mediated activation of receptor tyrosine kinases were down-regulated at the level of protein expression in the luminal lines relative to the basal lines. This suggests that in the luminal lines activities of kinases are less dependent on extracellular input than in the basal lines; this may be due to receptor overexpression in basal lines. We expected that subtypes would be more important in defining the signaling landscape, especially considering the large differences observed at the levels of protein, RNA expression, and genomic sequence. This lack of separation may reflect the targeted nature of our measurements or that there is actually a continuum of phenotypes rather than a clear luminal/basal separation.

The logic-based signaling network models that we built for each cell line served a double purpose. Firstly, model building condensed 1,995 points of information into 107 features, allowing visualization and enabling use of the models as predictors. Secondly, the models allowed the investigation of cell-line differences and provided insights into signaling mechanisms. We were surprised to find that the model parameters did not predict the responses to drugs, whereas the edge flux did mostly independent on the specific condition. This could be because the edge flux, maintains the correlation structure and although condition specific, integrates information about all conditions, and consequently information content does not change as a function of time or treatment as drastically as for the other predictors. The information transfer through the network as captured by the edge flux is important in defining a specific cellular state.

It was not surprising that the edge flux was not highly predictive for all drugs, since most drugs target processes not monitored by our antibody panel. The predictive accuracy of the edge flux was similar to or better than RNA-seq, despite of the almost 300-fold difference in the number of fitted features. Importantly, although we measured some cell-cycle markers, we did not model them. The edge flux was not predictive of sensitivities of drugs that interfere with the cell cycle, whereas median and cellular variability predictors were.

Using the edge flux, we were able to break down complex relationships between signaling and drug sensitivity. We showed that PKC is among the most important hubs, governing both sensitivity and resistance to many drugs. PKC is known to have complex functions in cancer progression (Garg et al., 2014). Interestingly, although PKC inputs (SERUM·PKC, EGFR·PKC, SRC·PKC, and BTK·SRC) were quite variable across cell lines, downstream activity, as assessed by model parameters, was similar across cell lines. Importantly, conservation at the model parameter level did not mean that the edge flux was highly conserved, as each edge flux strongly depended on all the upstream nodes. We also observed intriguing differential effects of ERK: The tumor suppressor activity of ERK was MEK dependent and involved 4EBP1 activation, whereas the tumor-promoting activity was FAK dependent and involved MSK1 and MSK2 activation.

Features correlating with sensitivity have potential in patient stratification. For example, cellular variability of p-STAT5 and p-STAT3 were predictive of lapatinib sensitivity and could therefore be used as a stratification marker. This is consistent with previous reports that inhibition of phosphorylation of JNK and STAT5 by lapatinib was observed only in sensitive cell lines (Gschwantler-Kaulich et al., 2016). *PI3K* mutational status is often not predictive of response to *PI3K*-targeted drugs. For example, only a subset of patients with mutant *PI3K* responded to pictilisib in clinical trials (Krop et al., 2016; Schmid et al., 2016; Schöffski et al., 2018). The edge flux and the median marker expression suggest important roles of p-MKK4 and p-STAT3 in defining pictilisib sensitivity; both have potential as markers for improving patient stratification.

Features correlating with resistance could aid in selection of efficacious combination therapies. Information on variability at the single-cell level allowed us to predict susceptibility to navitoclax, fulvestrant, NU7441, and AS601245, drugs for which other predictors were not informative. We observed a correlation between high EGFR- and RAS-independent activation of PI3K (SERUM·PI3K) to fulvestrant resistance, which suggests a potential combination therapy for fulvestrant resistance of fulvestrant with a PI3K inhibitor. Indeed, fulvestrant, which is used in endocrine therapy, was recently shown to be effective in postmenopausal women with endocrine-resistant, hormone receptor-positive, and HER2-negative advanced breast cancer when used in combination with a PI3K inhibitor (Baselga et al., 2017). Furthermore, the correlation of fulvestrant resistance to p-p38 and p-SRC cellular variability suggests that a combination of fulvestrant with either a p38 or a SRC inhibitor would be effective in treatment of ER-positive breast cancer. For lapatinib, the median expression of p-SRC was predictive of resistance; therefore, it would be interesting to test the effects of a combination therapy of lapatinib with a SRC inhibitor.

Our study thus makes a case for further expanding drug sensitivity predictors in the clinic with single-cell measurements, and we expect that a comprehensive signaling network model, which includes more markers covering more signaling pathways and integrates signaling single-cell heterogeneity, will further increase drug sensitivity prediction accuracy. Furthermore, the statistical power will improve with additional cell lines. Measurements of patient samples would be very difficult, firstly due to sample heterogeneity and secondly due to the complexity of cultivating patient-derived cells. Hence, translation of this knowledge to cheap, scalable, and robust biomarkers such as genomic signatures is needed before this type of modeling will impact patient care. As a proof-of-concept, we identified genomic aberrations that correlated with our drug-sensitivity predictive signaling features. For instance, through the lapatinib-predictive median expression levels of p-MKK3 and p-MKK6 we identified the known lapatinib sensitivity predictive *EP300* CNV (Mahmud et al., 2019) and the interesting *PPM1D* CNV, which encodes a serine threonine phosphatase amplified in approximately 8% of breast cancers (Lambros et al., 2010). Furthermore, the lapatinib-predictive p38·MK2 edge flux linked amplification of the breast cancer oncogene *A1CF* (Yan et al., 2017), and of *ASAH2B* to lapatinib resistance. Interestingly, the long non-coding *ASAH2B-2* was recently shown to promote breast cancer cell growth via the PI3K pathway (Li et al., 2018), supporting our findings. Furthermore, the fulvestrant sensitivity-predictive MSK·NF-κB edge flux could be linked to the breast-cancer proposed biomarker *MACC1* (Huang et al., 2013). Importantly, the same genomic aberrations were not directly predictive of drug sensitivity, showing the importance of the signaling characterization. Signaling characterization likely identifies non-linear relationships between drug sensitivity and the genomic aberrations. Our findings are consistent with the notion that multiple, potentially highly patient-specific mutations converge on common pathways. Thus, individual effects of mutations that we found to be associated with signaling features on drug sensitivity may have been too weak to yield statistically significant associations. However, multiple mutations together or different mutation in different patients may have modified the same pathways, which enabled us to detect pathway – drug sensitivity associations.

In summary, single-cell dynamic measurements of the cellular signaling response to stimulation and perturbation were used to construct logic-based mathematical models of cell signaling in cell lines from normal and cancerous breast tissue. The generated mechanistic signaling network models were predictive of resistance and sensitivity of cell lines to PI3K-MTOR and other drugs. Additionally, we identified genomic aberrations predictive of drug sensitivity though our signaling characterization. We envision that an approach similar to that introduced here will eventually deliver a robust drug-patient match.

## METHODS

### CELL CULTURE

Cells were obtained from suppliers or collaborators listed in Supplementary Table 4. With the exception of BT-474 cells, all were grown according to the supplier’s recommendation, and the maximum passage number was kept low (<15). Culture conditions are provided in Supplementary Table 5. All cells were free of *Mycoplasma*. For passaging, cells were incubated with 0.25% trypsin (Gibco) at 37 °C for 1 to 9 minutes depending on the cell line.

### STIMULATION

Cells were plated either in 150-mm or 100-mm dishes to achieve about 60% confluency at the time of analysis (maximum number of passages: 15). Cells were grown for 48 to 72 hours and then washed twice with PBS before starving them in serum-free medium without additives overnight before fixation for the time point zero profiling, stimulation, or treatment with inhibitor. For stimulation, EGF (Peprotech) and fetal bovine serum (FBS, Gibco) were added to final concentrations of 100 ng/ml and 10% v/v, respectively. For analysis of cells in the unstimulated state, starvation medium was replaced by complete medium. For experiments with inhibitor, the inhibitor was added 15 minutes before the addition of EGF and FBS. Inhibitors were diluted into starvation medium at approximately 100 fold the reported IC_50_ (Supplementary Table 3). At 1 hour before a time point, 5-iodo-2’-deoxyuridine was added to the medium at the final concentration of 4 μM. At 5 minutes before a time point, the dish was washed and then incubated with 2X TrypLE™ Express (Life Technologies) to induce cell detachment. At the time point, dishes were scraped and paraformaldehyde (PFA, from Electron Microscopy Sciences) was added to the cell suspension to 1.6% v/v, and cells were incubated at room temperature for 10 minutes. If EGF stimulation was not necessary, cells were harvested and crosslinked with PFA either immediately after or 15 minutes after inhibitor addition. PFA was then quenched with 40% w/v bovine serum albumin (BSA, Sigma) and after centrifugation, methanol chilled to −20 °C was used to resuspend the cells for long-term storage at −80 °C. Two individual experimental replicates (referred to as A and B) with partly overlapping time points (Supplementary Table 2) were performed for each cell line. For each replicate, the experimental procedures were performed on different days.

For the proteomic samples, cells were grown and collected in parallel with the replicates A and B. The third biological replicate (referred to as C) was grown independently and was passaged two more times to test for proteome stability over a limited number of cell divisions. Cells were plated in 150-mm dishes, grown for 48 to 72 hours, washed twice with PBS before addition of fresh complete growth medium overnight, washed, incubated with TrypLE™, and scraped. The sample was then washed with cold starvation medium, re-suspended in cold PBS, and sodium deoxycholate (DOC, from Sigma-Aldrich) was added to a final 5% w/w. The lysis was completed by on-ice sonication (2×30 seconds) and snap freezing in liquid nitrogen, before long-term storage at −80 °C.

### MASS CYTOMETRY SAMPLE PREPARATION AND MEASUREMENT

#### ANTIBODY CONJUGATION

The isotope-labeled antibodies (Supplementary Table 1) were generated using the MaxPAR antibody conjugation kit (Fluidigm) using the manufacturer’s standard protocol. After conjugation, antibody concentrations were determined based on absorbance at 280 nm. Candor PBS Antibody Stabilization solution was used to dilute antibodies prior to long-term storage at 4 °C. Antibody target specificity was previously confirmed (Lun et al., 2017) and optimal concentrations were determined by titration.

#### BARCODING AND STAINING PROTOCOL

The crosslinked and methanol-permeabilized cells were washed once with CSM (PBS with 0.5% w/v BSA, 2 mM EDTA) and once with PBS. Cells were incubated in PBS containing a barcoding reagent designed for a 126-well barcoding, containing four out of the nine metals used. Palladium (^105^Pd, ^106^Pd, ^108^Pd, ^110^Pd, Fluidigm) was used in conjunction with the chelating agent bromoacetamidobenzyl-EDTA (Dojindo); indium (^113^In, ^115^In, Fluidigm), yttrium, rhodium, and bismuth (^89^Y, ^103^Rh, ^209^Bi, Sigma Aldrich) were chelated to maleimido-mono-amide-DOTA (Macrocyclics). The samples were randomly distributed across the wells and incubated for 30 minutes at room temperature at 100 nM metal concentrations except for bismuth (20 nM), indium isotope 113 (200 nM), and rhodium (2 μM). After incubation, the sample was washed four times with CSM (Bodenmiller et al., 2012), pooled, and stained with the metal-conjugated antibody mix at 4 °C for 1 hour. The antibody mix was removed by washing cells three times with CSM. For DNA staining, iridium-containing nucleic acid intercalator (^191^Ir and ^193^Ir, Fluidigm) diluted in PBS with 1.6% PFA was added to the cells to a final concentration of 500 µM, and cells were incubated at 4 °C overnight. The following day, the intercalator solution was removed, cells washed sequentially with CSM, PBS, and ddH_2_O and stained for cell volume with 12.5 μg/ml bis(2,2’-bipyridine)-4’-methyl-4-carboxybipyridine-ruthenium-N-succidimyl ester-bis(hexafluorophosphate) (^96^Ru, ^98-102^Ru, ^104^Ru, Sigma Aldrich) in 0.1 M sodium hydrogen carbonate, pH 8.3 (Sigma Aldrich) for 10 minutes at room temperature. Subsequently, samples were washed once each with CSM, PBS, and ddH_2_O. After the last washing step, cells were resuspended in cell running buffer (Fluidigm) and EQ^TM^ Four Element Calibration Beads (Fluidigm) were added in a 1:10 ratio (v/v). Subsequently, samples were filtered through a 35-μm strainer just before the mass-cytometry measurement.

#### MASS CYTOMETRY ANALYSIS

Samples were analyzed on an upgraded CyTOF2 (Fluidigm) using the Super Sampler (Victorian Airship) introduction system. The manufacturer’s standard operation procedures were used for acquisition at a cell rate of ∼300 cells per second. After the acquisition, all FCS files from the same barcoded sample were concatenated as previously described (Bodenmiller et al., 2012), data were then normalized, and bead events were removed (Finck et al., 2013). A doublet-filtering scheme and single-cell deconvolution algorithm (Zunder et al., 2015) were used to achieve doublet removal and for de-barcoding of cells into their corresponding wells. Subsequently, data were processed using Cytobank (Kotecha et al., 2010). Additional gating on the DNA channels (^191^Ir and ^193^Ir) was used to remove remained doublets, debris, and contaminating particulates. FCS files were exported and loaded into R for downstream analysis (*flowCore* Bioconductor/R-package).

Firstly, compensation for channel crosstalk was performed using single-stained polystyrene beads (Chevrier et al., 2018). Secondly, samples that were measured multiple times were combined and medians and the coefficients of variation (cellular variability) were computed. Thirdly, median signal intensities per channel were arcsinh-transformed (asinh(x+1), *flowCore* Bioconductor/R-package). Fourthly, to control for batch effects, all cell lines were processed, stained, and measured together with five samples that served as technical replicates. The technical replicates were generated in large batches from two cell lines (HCC70 and MDA-MD-453) prepared at different time points to ensure negative and positive controls for each measured marker. The simultaneous processing enabled direct quantitative comparisons within a cell line and the technical replicates enabled identification of batch effects and were used in batch-correction quality control. Both the t-SNE algorithm (Van Der Maaten and Hinton, 2008) (*Rtsne* R-package) and principal component analysis (PCA, *sparsepca* R-package) were used for identification of batch effects. While the cellular variability showed no detectable batch effect, the median showed an overall loss in signal strength between the first fifteen measured cell lines and the rest (likely due to antibody aging). Centering the two batches to the common average (μ_total_) for each median value (x_centered_ = x - μ_batch_ + μ_total_) prevented the formation of batch-specific clusters by both PCA and t-SNE. Fifthly, the biological replicates (time course A and B, which had only partially overlapping time points) were integrated into one consensus measurement. The two time courses were centered by subtracting the means of the individual time courses and addition of the overall mean. Finally, the unmeasured time points were linearly interpolated using the R function *approxfun* (the same process was used for missing or slightly different time points) and the averages between the two biological replicates were computed. For the time point 60 minutes, sampled only in time course B, the value for the time course B was taken directly without extrapolating a value for the second time course.

#### IDENTIFICATION OF GENERAL TRENDS

To characterize the general signaling response across all 67 cell lines we performed a two-way ANOVA comparing treatment and time (with an interaction term). The obtained p-values were corrected for multiple hypothesis testing using the Benjamini and Yekutieli multiple hypothesis correction, and, if relevant, the significant relationships (FDR 5%) were further characterized using Tukey honest significant differences computation. The increase in heterogeneity following inhibition was quantified by comparing the average standard deviation of the median fold change at 60 minutes with ANOVA followed by Benjamini and Yekutieli multiple hypothesis correction for the six treatments.

### MASS SPECTROMETRY SAMPLE PREPARATION AND MEASUREMENT

#### DIGESTION

Cell lysates from independent biological replicates were aliquoted in equivalent volumes containing 100 μg of proteome sample (quantified with a BCA assay). The samples were then reduced with 5 mM Tris(2-carboxyethyl)phosphine (ThermoFisher Scientific) for 30 minutes at 37 °C and then alkylated in the dark for 30 minutes at 25 °C with 40 mM iodoacetamide (Sigma Aldrich). Samples were diluted with 0.1 M ammonium bicarbonate (Sigma Aldrich) to a final concentration of 1% DOC before overnight digestion at 37 °C with lysyl endopeptidase (Wako Chemicals) and sequencing-grade porcine trypsin (Promega) at an enzyme-substrate ratio of 1:100 for both. Trypsin was inactivated by adding formic acid (AppliChem) to a final concentration of 1% v/v, and the precipitated DOC was removed by centrifugation. The acidified peptide mixtures were loaded into 96-well elution plates (Waters), desalted, and eluted with 80% acetonitrile. Samples were dried in a vacuum centrifuge, solubilized in 0.1% formic acid, and analyzed by mass spectrometry.

#### SPECTRAL LIBRARY SAMPLE PREPARATION

The spectral libraries were obtained from all cell lines with at least three biological replicates (12 cell lines) or with eight fractions from two independent cell line pools. The cell lines pools were composed of 27 and 28 cell lines, respectively. The four central fractions were measured twice. The fractionation was performed using the Pierce™ high pH, reversed-phase peptide fractionation kit (Thermo Fisher Scientific), and iRT peptides (Byognosis AG) were added.

#### PEPTIDE SEPARATION

Digested samples were analyzed on an Orbitrap Q Exactive Plus mass spectrometer (Thermo Fisher Scientific) equipped with a nano-electrospray ion source and a nano-flow LC system (Easy-nLC 1000, Thermo Fisher Scientific). Peptide separation was performed on a 40 cm x 0.75 μm i.d. column (New Objective, PF360-75-10-N-5) in-house packed with 1.9-µm C18 beads (Dr. Maisch Reprosil-Pur 120) and heated to 50 °C. Buffer A was 0.1% w/v formic acid, and buffer B was 0.1% w/v formic acid in acetonitrile. The flow rate was 300 nL/min. The gradient was as follows (buffer B in buffer A): linear from 5% to 25% over 100 minutes, 25% to 40% over 10 minutes, and 40% to 90% over 5 minutes, finishing with isocratic 90% for 5 minutes.

#### SPECTRAL LIBRARY DATA-DEPENDENT ACQUISITION

For the spectral library, the 55 samples were analyzed by shotgun LC-MS/MS data dependent acquisition (DDA) by injection of 1 μL peptide digests at a concentration of 1 μg/μL. The MS1 spectra were acquired from 350 to 1,500 m/z at a resolution of 70,000, and the 20 most intense precursors exceeding 1,300 ion counts were selected for fragmentation at 25 eV normalized collision energy. The MS2 spectra were acquired at a resolution of 17,500 with maximally 100,000 ions, collected for 55 ms maximally. All multiply charged ions triggered MS-MS scans followed by a 30-second dynamic exclusion, and singly charged precursor ions and ions of undefinable charged states were excluded from fragmentation.

#### PEPTIDE IDENTIFICATION AND SPECTRAL LIBRARY GENERATION

The DDA spectra were searched against the canonical Human Uniprot fasta database (version August 2018) using the Sequest HT database search engine in Protein Discoverer (version 2.2.0.388, Thermo Fisher Scientific). We allowed for up to two missed cleavages, excluded cleavage of KP and RP peptide bonds and applied a full tryptic digestion rule. Cysteine carboxyamidomethylation (+57.021 Da) and methionine oxidation (+15.995) were allowed as static and dynamic modifications, respectively. Monoisotopic peptide tolerance was set to 10 ppm, and fragment mass tolerance to 0.02 Da. The identified proteins were filtered using the high peptide confidence setting (1% false discovery rate (FDR) on peptide level). For generation of the spectral library the DDA spectra analyzed as described above were imported in the software Spectronaut Pulsar (11.0.18108.11.30271 Asimov, Biognosys AG) (Bruderer et al., 2015).

#### DATA-INDEPENDENT ACQUISITION

After addition of iRT peptides (Biognosys AG), 1 μL of peptide digest from each biological replicate was injected independently at a concentration of 1 μg/μL and measured in data-independent acquisition (DIA) mode. The DIA-MS method was as previously described (Piazza et al., 2018). Briefly, an MS1 survey scan was performed from 350 to 1,500 m/z at a resolution of 70,000 with AGC target of 3×10^6^ and a 120-ms injection time. The twenty variable-width windows optimized to equally distribute the number of precursor ions had a 1 m/z overlap. MS2 spectra were acquired at a resolution of 35,000 with a fixed first mass of 150 m/z and an AGC target of 1×10^6^. In order to mimic DDA fragmentation, the normalized collision energy was 25 eV based on the doubly charged center m/z of the isolation window. The maximum injection times were automatically chosen to maximize parallelization resulting in an approximate 3-second duty cycle.

#### DIA-MS TARGETED DATA EXTRACTION

Targeted data extractions of DIA-MS acquisitions were performed with Spectronaut Pulsar (11.0.18108.11.30271 Asimov, Biognosys AG) with default settings (Bruderer et al., 2015). Retention time prediction was set to dynamic with correction factor 1 for XIC extraction window determination. Non-linear calibration was used for retention time correction, and MS2 level interference correction was enabled (Bilbao et al., 2015). The FDR was set to 1% at peptide precursor level and was estimated with mProphet (Reiter et al., 2011). The method compares time-resolved MS/MS maps measured with the DIA-MS method as previously described (Gillet et al., 2012).

#### QUANTIFICATION

Subsequently the data were analyzed with *MSstats* (using Tukey’s median polish, R package version 3.10.6) for differential protein abundance in comparison to five cell lines derived from normal breast tissue (184A1, 184B5, MCF 10A, MCF 10F, and MCF 12A) (Choi et al., 2014). For quantification, only proteotypic peptides, which are uniquely present in the sequence of one protein, were used. The default settings for *SpectronauttoMSstatsFormat* and the *dataProcess* functions were used with the exception that the normalization was provided by Spectronaut Pulsar and zero intensities were censored. For each cell line comparison, MSstats (with the *groupComparison* function) was used to determine model-based estimates of fold changes. The adjusted p-values were determined using the Benjamini-Hochberg method to control the FDR at the cut-off level of 0.05 (Benjamini and Hochberg, 1995). Proteins with a fold change of at least 2 were considered as differentially abundant. For plotting, *-Inf*, *NA*, and *Inf* values were set to the dataset’s minimum, zero, and maximum, respectively.

#### BREAST CANCER-ASSOCIATED PROTEINS

The list of curated breast carcinoma-related genes was downloaded from the human disease discovery platform Disgenet (http://www.disgenet.org/, *curated_gene_disease_associations.tsv* file, downloaded on April 12, 2018) (Piñero et al., 2017). We included in the list genes annotated to the following breast-cancer disease categories: ‘Malignant neoplasm of breast’, ‘Breast Cancer, Familial’, ‘Hereditary Breast and Ovarian Cancer Syndrome’, ‘Breast Carcinoma’, ‘Breast Neoplasms, Male’, ‘Invasive Ductal Breast Carcinoma’, ‘Inflammatory Breast Carcinoma’, and ‘Breast Diseases’. The list includes a total of 53 genes among which 38 are associated with significant changes in protein abundance in one or multiple cell lines.

#### GENE ONTOLOGY ENRICHMENT ANALYSIS

The Gene Ontology (GO) enrichment analysis was performed using the GOATOOLS Python-based library (Klopfenstein et al., 2018). The background set corresponds to all human proteins (https://www.ncbi.nlm.nih.gov/genome/51, genome annotation file downloaded on May 12, 2015). The option *propagate_counts* was set to *False* to avoid propagation of the annotations of a gene from the assigned GO category to all parent GO terms. The p-value was calculated using Fisher’s exact test and then adjusted for multiple testing using the Benjamin-Hochberg correction method (Benjamini and Hochberg, 1995). We next calculated the specificity of the enriched GO terms by computing their information content (IC) as follows: IC = −*log* (frequency), where frequency is the number of genes annotated to the current GO term divided by the total number of associations between genes and GO terms in the full branch. The semantic similarity between all the enriched GO terms was then calculated as the inverse of the semantic distance (number of branches separating the terms). The IC and semantic similarity values were finally used to filter the list of GO-enriched categories selecting the more informative GO term (highest IC) among pairs of terms showing a semantic similarity higher than 0.5. To visualize the results of the functional enrichment analysis across the 62 tumor-derived cell lines, GO terms with a number of associated genes larger than 500 or smaller than 100 as well as terms commonly enriched in less than 10 cell lines were excluded to reduce the complexity and the redundancy of the plot while preserving the biological outcome.

### SIGNALING MODEL

We utilized the logic-based ordinary differential equation (ODE) formalism (Terfve et al., 2012) to model the signaling network of each cell line. This is a semi-mechanistic approach that combines perturbation data with prior knowledge, such as protein-protein interactions. The goal was to describe the signal transduction upon perturbation through protein activation cascades.

#### LOGIC-BASED DYNAMIC MODELS

First, we built a prior knowledge network (PKN) that contains nodes (protein markers); signed, directed edges (interactions of the proteins from Omnipath); and logical gates (AND, OR) reported in a simple interaction file (SIF) that contains three columns: source node, interaction type (activation or inhibition), and target node (Türei et al., 2016) (Sup. Fig. 3A and Sup. Table 6). This PKN was built around the measured and perturbed proteins in the experiments and was used for all cell lines. Next, the PKN was translated into a dynamic ODE model using the CNORode modeling package (https://github.com/saezlab/CNORode) that is part of the CellNOpt family (Terfve et al., 2012). The node *i* property *x_i_* ∈ [0,1], *i* = 1 … *N* in the network, the differential equation was written as in Equation 1.

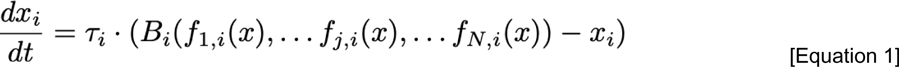

τ*_i_* is the responsiveness parameter of the node *x_i_*, and a larger value results in a faster response to change in the node. *B_i_* is the continuous Boolean function: *B_i_* : [0,1]*^N^* -> [0,1]. This function accounts for the AND and OR gates of the incoming edges on node *i* (Wittmann et al., 2009). *f_ij_(x)* is the transfer function from node *j* to node *i*; it describes how node *i* depends on node *j*, here we use a version of the previously described Hill-type function (Eduati et al., 2017) as shown in Equation 2.

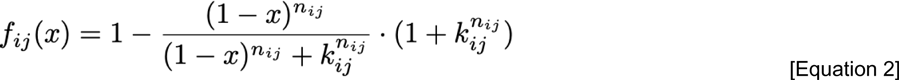

This function has a sigmoidal-shape characterized by the free parameters *κ_ij_* and *n_ij_*. The trajectories are constrained in the [0-1] interval. The extreme case of 0 means that the corresponding node is inactive or inhibited, and 1 means that the node is fully activated. In order to compare the experimental measurements with the simulation results, the data were scaled to the [0-1] interval. Each marker was scaled separately across all cell lines and conditions, but different markers were not compared due to differences in sensitivity. First, the median data were scaled to the 0-1 range using the 99% interquartiles, as described in Equation 3.

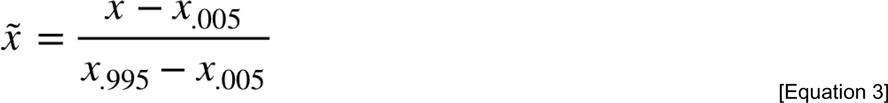

Then values >1 or <0 were set to 1 and 0, respectively. For each experimental condition, the corresponding states in the model were adjusted. For instance, application of the stimulation was modeled by setting the input nodes to 1, and inhibition was modeled by setting the inhibited nodes to 0.

#### CELL LINE-SPECIFIC PARAMETER ESTIMATION

All the cell-line models were built from the same PKN, and 67 cell line-specific ODE models were generated. In these models there were four types of unknown parameters:

1. Initial conditions for unmeasured states, totaling 11 parameters
2. A node responsiveness parameter for each node (τ*_i_*), totaling 40 parameters
3. The edge parameter *n*, totaling 88 parameters
4. The edge parameter κ, totaling 88 parameters

In order to reduce complexity, we fixed the values of the *n* parameters to 3, since they influence the outputs the least. Then we trained the ODE models using the CNORode package, following the standard approach with minor improvements. In short, the package relies on the MEIGOR optimization toolbox to find the parameter values that results in the best fit (measured by the root mean squared error, RMSE) (Terfve et al., 2012). We applied L2 regularization in the optimization as previously introduced to the CNORode (Eduati et al., 2017) to cope with non-identifiability of estimated parameters and to reduce overfitting. We evaluated five cell-line models for tuning the regularization parameter. The value of 1×10^-5^ for the regularization parameter resulted in a good balance between sparsity and fit (data not shown). Each model was trained using the global optimiser enhanced Scatter Search (eSS) together with the Dynamic Hill Climbing local search algorithm with 10 optimizations, 20 minutes each (as implemented in the MEIGOR package). After the optimization, the model simulation was plotted against the data to evaluate the fitting quality. To control for the model quality both the *r^2^* as well as the root mean square error (RMSE) were evaluated. The biological variance was computed as the RMSE between the median and the single biological replicates. The random model was made the same way as the cell line models for all cell lines, but parameters were randomly generated and not optimized.

#### MUTATION MATRIX

To determine the mutational status, the SNP-specific genotype as defined by the Illumina HumanOmni1-Quad v1.0 Multi-UseManifest File (http://emea.support.illumina.com/downloads/humanomni1-quadv1_mu_product_files.html, downloaded on July 18, 2019) was mapped to the NCBI 37.1 (GRCh37) genome build taking into account ambiguous IUPAC notations and indel information. Subsequently, a cell line × SNP mutation matrix was constructed, where elements denote the fraction of alleles for which the cell line genotype mismatches the reference genotype for the respective SNP. Finally, two gene-level mutation matrices were obtained by first mapping the SNP entries to genes using the *pyensembl* package (Ensembl release 55) and then counting the values exceeding the threshold *q* (*q* = 0.5 for a “dominant” matrix and *q* = 1 for a “recessive” one) for each cell line and gene.

### DRUG SENSITIVITY PREDICTION

We predicted the response (-ln IC_50_) using Macau for each drug (409 vectors of -ln IC_50_ values for 347 individual compounds) in 48 cell lines (training) with data available in the GDSC dataset (Simm et al., 2017; Yang et al., 2013). A 5-fold cross-validation and 40 iterations were used to obtain an average prediction performance (performance score) and an adjusted *p*-value for each predictor (Benjamini and Hochberg, 1995). As input for the drug response prediction we selected the RNA-seq filtered for exons (Marcotte et al., 2016), the protein abundance ratio, the estimated parameters (model parameters), the single-cell coefficient of variation (cellular variability), the median marker expression (median), and the edge flux.

#### EDGE FLUX

The edge fluxes were computed as the time-dependent edge strengths, or in other words the transfer function from node *j* to node *i* determined as describe by Equation 2, for each specific condition (treatment and time). This equation describes how much a node is influenced by its upstream nodes at a certain time and treatment condition. The edge flux value depends on the model parameters as well as on the activity of the influencing/upstream nodes (*x*, Equation 3) and can be seen as the signaling flux of the edges at a specific time and treatment condition. For the *AND*-gated edges, the edge flux is the product of two *f*-functions (Equation 2). For instance, AKT,RAS·MEK^S221^ integrates positive RAS as well as negative AKT influence and is therefore influenced by both *f_AKT_(x_AKT_)* and *f_RAS_(x_RAS_)* as follows: AKT,RAS·MEK^S221^ = *(1-f_AKT_) * f_RAS_*.

#### MACAU

Macau is a matrix factorization algorithm based on Markov chain Monte Carlo (MCMC) sampling that incorporates information in rows and/or columns to improve the accuracy of the predictions (Simm et al., 2017). Cell line-specific information was transformed into a matrix of L latent dimension (set to 10 as we only used cell line features) by a link matrix. Drug response was then computed by a matrix multiplication of the two latent matrices, from drug (L_drug_) and cell lines (L_cell_) sides. Macau employs Gibbs sampling to sample both the latent vectors and the link matrix, which connects the additional information to the latent vectors. For MCMC sampling, we chose a burn in of 400 samples, then collected 600 samples. After collection of each sample, we predicted drug response by multiplying the two latent matrices and then averaged across all 600 samples. We used a 5-fold cross-validation and iterated 40 times for an average prediction performance (performance score) (Simm et al., 2017; Yang et al., 2018). For prediction of responses of cells lines not part of the training exercise, we used the same strategy except that inside the cross validation loop we predicted not on the test set but on the 19 hold-out cell lines.

#### FEATURE IMPORTANCE

The procedure for estimating the feature importance was as follows:

1. For every MCMC sample, we:

a. Extracted the latent vector V_drug 1_ for a given drug number 1. This latent vector represented a subset of L_drug_ for a specific drug 1.
b. Extracted the link matrix of the cell-line side, Link_cell_.
c. Computed the element-wise multiplication: V_drug 1_ **\*** Link_cell_. The resulting matrix represents the feature importance of the predictors for each latent dimension.
2. For each predictor, averaged across all L latent dimensions, we:

a. Took the average feature importance across all cross validations and over 40 iterations.
b. Generated random permutations of the feature importance matrix for all drugs 1,000 times, where we shuffled the predictors for each cell line independently. We then derived an empirical null distribution for each feature importance value. If the value was positive, we defined the p-value as the number of cases in the null distribution greater than the value of interest divided by the number of permutations. If the value was negative, we defined the p-value as the number of cases in the null distribution smaller than the value of interest divided by the number of permutations.

#### FEATURES PREDICTIVE OF OVERALL SENSITIVITY

To determine features that significantly predicted overall sensitivity we performed an ANOVA with H_0_: effect size > 0.01 or < −0.01 across all drugs and conditions for the predictors: median, cellular variability, and edge flux. The obtained p-values were corrected for multiple hypothesis using Benjamini and Yekutieli (features * 2).

#### FEATURES PREDICTIVE OF PATHWAY SPECIFIC SENSITIVITY

Drugs were binned in different classes according to the target pathway (Yang et al., 2013). Upon selection of ten interesting classes we looked for features predictive for these specific dug classes with an ANOVA with H_0_: effect size > 0.01 or < −0.01. The obtained p-values were corrected for multiple hypothesis testing using Benjamini and Yekutieli (features * 2), and the significant relationships (FDR 5%) were further dissected using Tukey honest significant differences computation.

### IDENTIFICATION OF PREDICTIVE GENOMIC ABERRATIONS

#### RANDOM FOREST QTL MAPPING

Mapping of quantitative trait loci (QTLs) was conducted using random forest ensemble learning (Michaelson et al., 2009), an approach that has been shown to outperform legacy and other multi-locus QTL mapping methods (Michaelson et al., 2010). Briefly, a random forest classifier was trained on genetic markers to predict relevant signaling features, and resulting marker selection frequencies were used as a measure for the strength of the respective QTL. Separate analyses were conducted for genotype matrices assembled from publicly available SNP (for 61 cell lines) and CNV (for 64 cell lines) data obtained using Illumina HumanOmni1-Quad BeadChips (Marcotte et al., 2016). Gene-level CNV data was discretized by thresholding CBS-segmented log-R ratios at −0.2 and 0.2. Genetic markers were filtered for mutations in cancer-driver genes (Forbes et al., 2011; Sondka et al., 2018). Markers with missing values for more than half of the cell lines and markers with a major allele frequency larger than 0.9 were excluded, resulting in 736 SNPs and 518 CNVs. The proportion of genotypic variance included in the estimated population-structure covariates was set to 0.75. Mapping scores were computed by training 10 random forests (representing independent imputations of missing marker values), each comprising 1,000 trees. To determine the significance of identified QTLs, an empirical null distribution of marker selection frequencies was estimated by repeating this step 5,000 times for 100 independent phenotype vector shuffles, totaling 5 billion randomized trees per analyzed experimental condition. The product of zero-clipped permutation importance (PI) and average increase in node purity (RSS), normalized across traits, was used as a readout for feature importance. Finally, empirical p-values adjusted for multiple testing were computed and checked for convergence by plotting them against the number of batches at varying intervals (Benjamini and Hochberg, 1995). All computations were conducted using modified versions of the *RFQTL* and *RandomForest* R packages and executed in parallelized fashion on local cloud infrastructure (Michaelson et al., 2010). Due to the small dataset used we selected a permissive adjusted *p*-value cutoff of 0.3. The same method, cutoffs, and selection of genomic aberrations were used for a QTL analysis on the IC_50_ values (*–ln*) for pictilisib, fulvestrant, and lapatinib in 42 and 40 cell lines for the CNVs and SNPs, respectively.

#### COMPARISON OF THE TWO QTL APPROACHES

The genomic features (independently for SNP and CNV) passing the 0.3 adjusted *p*-value cutoff were subsequently tested for effects on the drug sensitivity (IC_50_ values (*–ln*) for pictilisib, fulvestrant, and lapatinib) by using ANOVA. The ANOVA was performed only when the genomic feature identified by QTL was linked to a signaling feature predictive of the sensitivity of the specified drug (p-value ≤ 0.005 and effect size ≥ ± 0.02, see the section “Feature Importance”). The p-values obtained by the ANOVA were then corrected for multiple hypothesis testing for each drug individually (Benjamini and Hochberg, 1995).

## ACKNOWLEDGEMENTS

We would like to thank the Bodenmiller and Picotti laboratories for support and fruitful discussions. In particular, we would like to thank Andrea Jacobs, Stefanie Engler, and Paul Boersema for their help in running the labs. We would especially like to thank Prof. Benjamin G. Neel, NYU Langone Health, and Prof. Joe Gray, Oregon Health & Sciences University Center for Spatial Systems Biomedicine, Portland, for providing cell lines. We would like to thank Jan Großbach (CECAD, University of Cologne, Germany) for support with the random forest QTL mapping. B.B. was supported by a SNSF R’Equip grant, a SNSF Assistant Professorship grant, the SystemsX Transfer Project “Friends and Foes”, the SystemX grants Metastasix and PhosphoNEtX, a NIH grant (UC4 DK108132), the CRUK IMAXT Grand Challenge, and by the European Research Council (ERC) under the European Union’s Seventh Framework Program (FP/2007-2013)/ERC Grant Agreement n. 336921. J.S.R. was partially funded by the Joint Research Center for Computational Biomedicine (which is partially funded by Bayer AG). A.B. received funds from the German Federal Ministry of Education and Research (BMBF grant: PhosphoNetPPM).

## AUTHOR CONTRIBUTIONS

MT, AB, PP, JSR, and BB conceived and designed the experiments. MT performed experiments. VC and MT analyzed the proteomic data. Mass cytometry data pre-processing was performed by KC and MT, while AG build the logic models with the help of MT. Drug sensitivity prediction was done by MY with help by AG and MT. The genomic analysis was performed by JW and MT. MT, NdS, and BB drafted the manuscript. All authors contributed to the manuscript.

## DATA AND SOFTWARE AVAILABILITY

The mass spectrometry proteomics data have been deposited to the ProteomeXchange Consortium via the PRIDE partner repository with the dataset identifier PXD017199. The mass cytometry data and other analyses including all cell-line-specific logic-based models and drug sensitivity predictions will be available through Sage and Mendeley Data upon final publication. All codes are provided in Github (https://github.com/BodenmillerGroup/Signaling-Network-Landscape-of-Breast-Cancer) and will be updated until final publication.

---

**Supplementary Figure 1:**
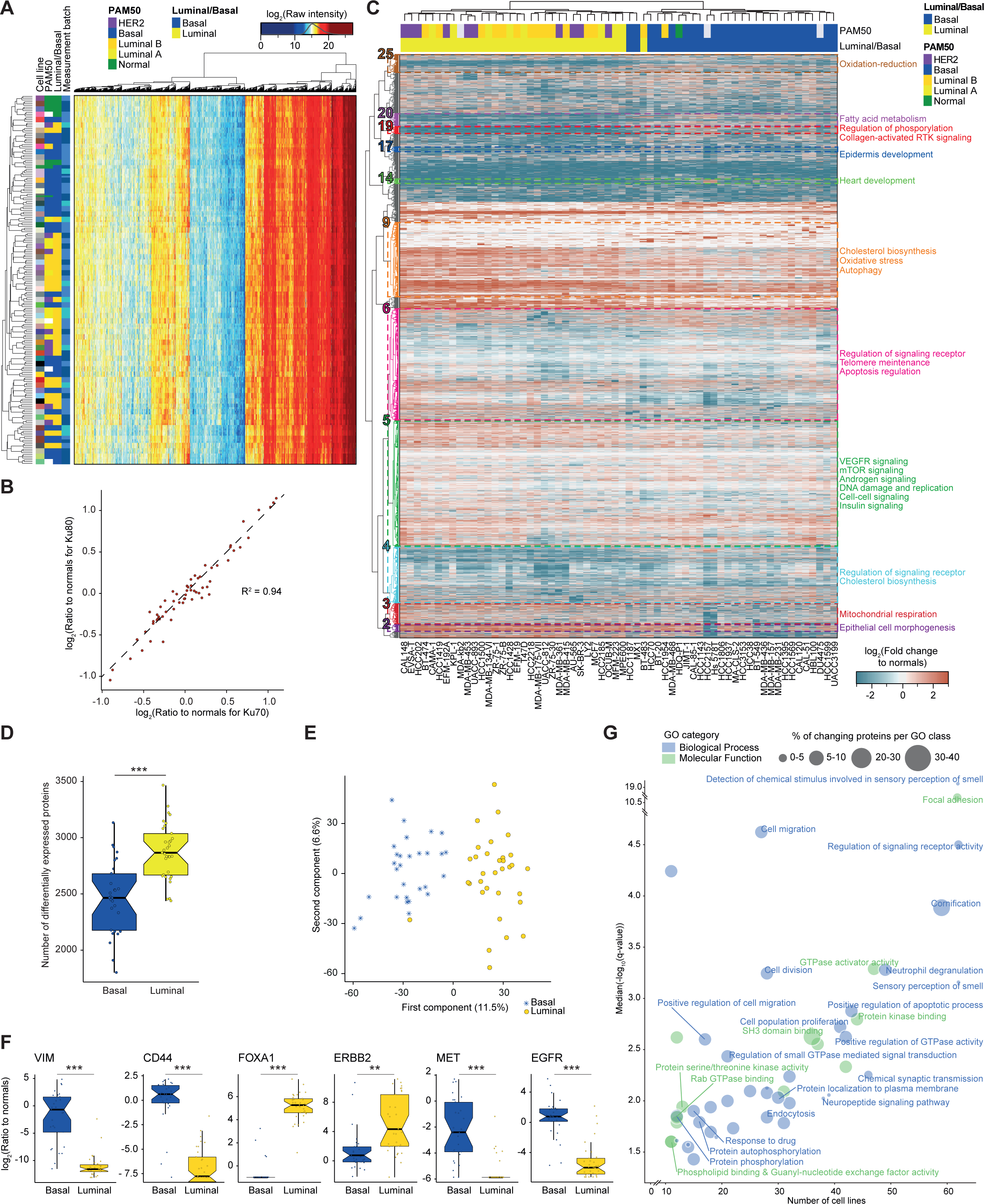
The Proteomes of Breast Cancer and Normal Breast Cell Lines. *(A) Clustering of the measured samples based on the 2,762 consistently quantified proteins. Color intensity on the white-red scale indicates log_2_ raw protein abundance values. The prediction analysis of microarray 50 (PAM50) classifications of each cell line, the curated luminal/basal classifications, cell line, and the measurement batch are shown. The data represent three independent experiments*. *(B) Values of Ku70 abundance plotted versus Ku80 abundance for all cancer cell lines. The function y = x is indicated by a dashed line and the coefficient of determination is reported*. *(C) Clustering of the cell lines based on protein levels. Color intensity on the blue-to-red scale indicates log_2_ protein abundance ratio in cancer cell lines relative to normal cell lines. Hierarchical clustering of the cell lines based on the 9,031 identified proteins and GO enrichment categories of the 24 identified clusters are shown. Clusters with significant enrichment (adjusted p-value < 0.05) for at least one biological process are highlighted in different colors, and significant terms are summarized. The prediction analysis of microarray 50 (PAM50) classifications of each cell line and the curated luminal/basal classifications are also shown*. *(D) Number of differentially expressed proteins plotted against luminal/basal classification (33 and 29 cell lines, respectively). Thick lines indicate medians, boxes indicate the 25% and 75% quantiles, whiskers extend between the median and ± (1.58 * inter-quantile range). Each data point represents a cell line (ANOVA, ***p ≤ 0.001)*. *(E) Principle component analysis representation of the log_2_ protein abundance along the first two components (sparsepca R-package). The data for luminal and basal cell lines (33 and 29 cell lines, respectively) are blue asterisks and yellow points, respectively. The first component separates cell lines by classification and explains 11.5% of the variance in the data*. *(F) Protein abundances for VIM, CD44, FOXA1, ERBB2, MET, and EGFR plotted against luminal/basal classification (33 and 29 cell lines, respectively). These proteins are six of the 1,946 significantly differentially expressed proteins (ANOVA with multiple hypothesis correction for the number of quantified proteins with Benjamini and Yekutieli: *p ≤ 0.05, **p ≤ 0.01, ***p ≤ 0.001). Thick lines indicate medians, boxes indicate the 25% and 75% quantiles, whiskers extend between the median and ± (1.58 * inter-quantile range). Each data point corresponds to a cell line*. *(G) Median q-values for the enriched GO-terms plotted against the number of cell lines in which the GO term is significant. Bubble size is proportional to the median percentage of proteins changing within the GO term and in blue and green are the biological process and molecular function categories, respectively*.

**Supplementary Figure 2:**
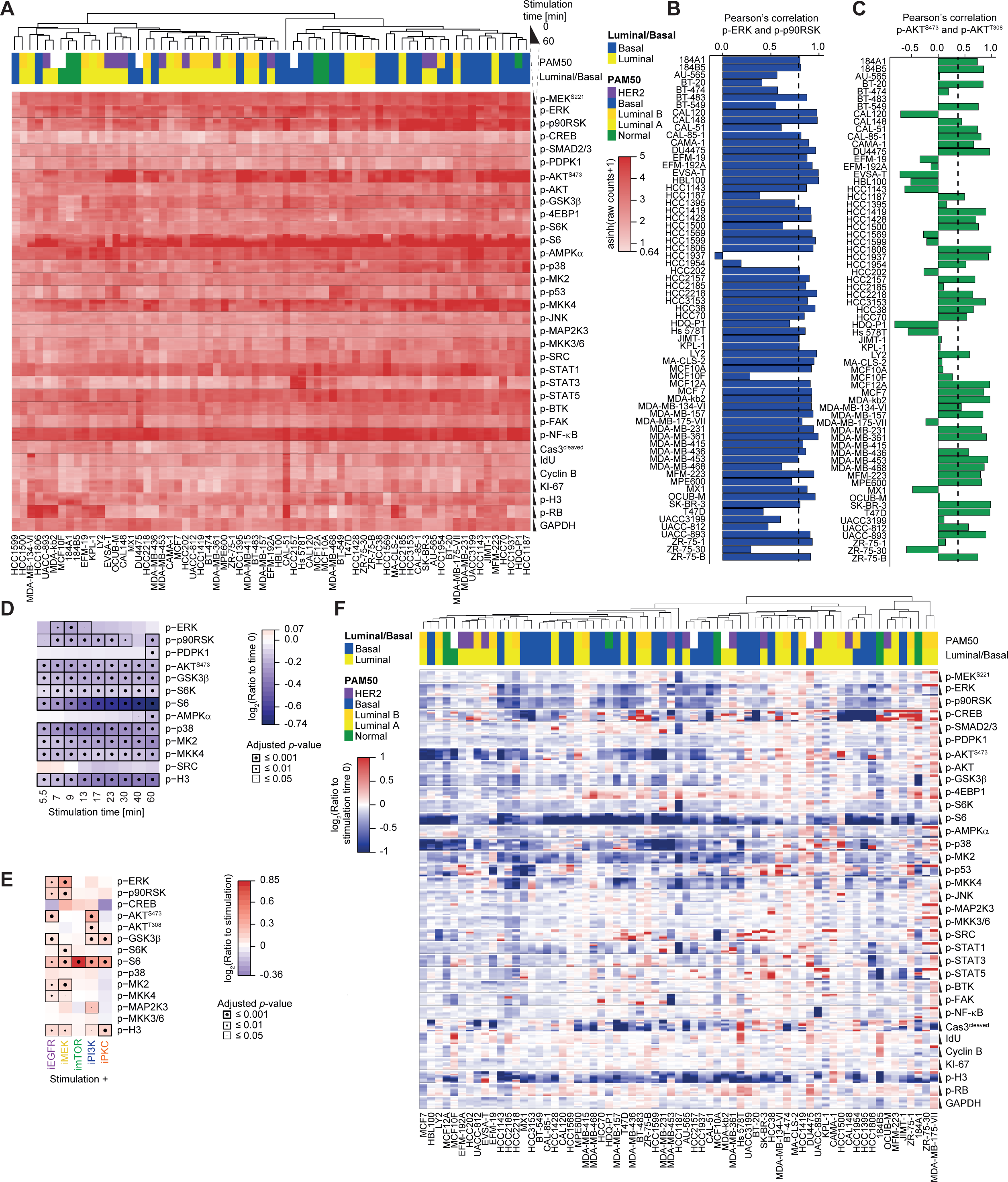
The Signaling Landscape of Breast Cancer Cell Lines. *(A) Heat map of marker abundances, ordered by increasing stimulation time and treatment, in all 67 cell lines. The cell lines were clustered based on their signaling signature. PAM50 tumor subtype classification and luminal/basal classification are overlaid. Data are from two independent experiments combined by linear interpolation*. *(B) Pearson’s correlation between p-ERK and p-p90RSK ratios to EGF stimulation time zero across the 67 cell lines. A vertical dashed line indicates the median. p-ERK and p-p90RSK are usually highly correlated, presumably due to the underlying regulatory mechanisms; however, in some cell lines this relationship is not observed and in HCC1937 cell line abundances are anti-correlated*. *(C) Pearson’s correlation between the ratio of the two AKT phosphorylation sites (p-AKT^S473^ and p-AKT^T308^) and EGF stimulation time zero across the 67 cell lines. A vertical dashed line indicates the median. Strikingly, in some lines the abundances are strongly anti-correlated, as in HDQ-P1 and EVSA-T*. *(D) Ratios of single-cell coefficient of variation of markers with significant differences over time in all 67 cell lines. The adjusted p-value is indicated by the dot size and the box thickness*. *(E) Ratios of single-cell coefficient of variation of markers with significant effects of the indicated kinase inhibitors compared to EGF stimulation alone. The adjusted p-value is indicated by the dot size and the box thickness*. *(F) Ratios of single-cell coefficient of variation to EGF stimulation time zero, ordered by increasing stimulation time and treatment for the 67 cell lines clustered based on their signaling signatures. PAM50 tumor subtype classification and luminal/basal classification are overlaid. Data are from two independent experiments combined by linear interpolation*.

**Supplementary Figure 3:**
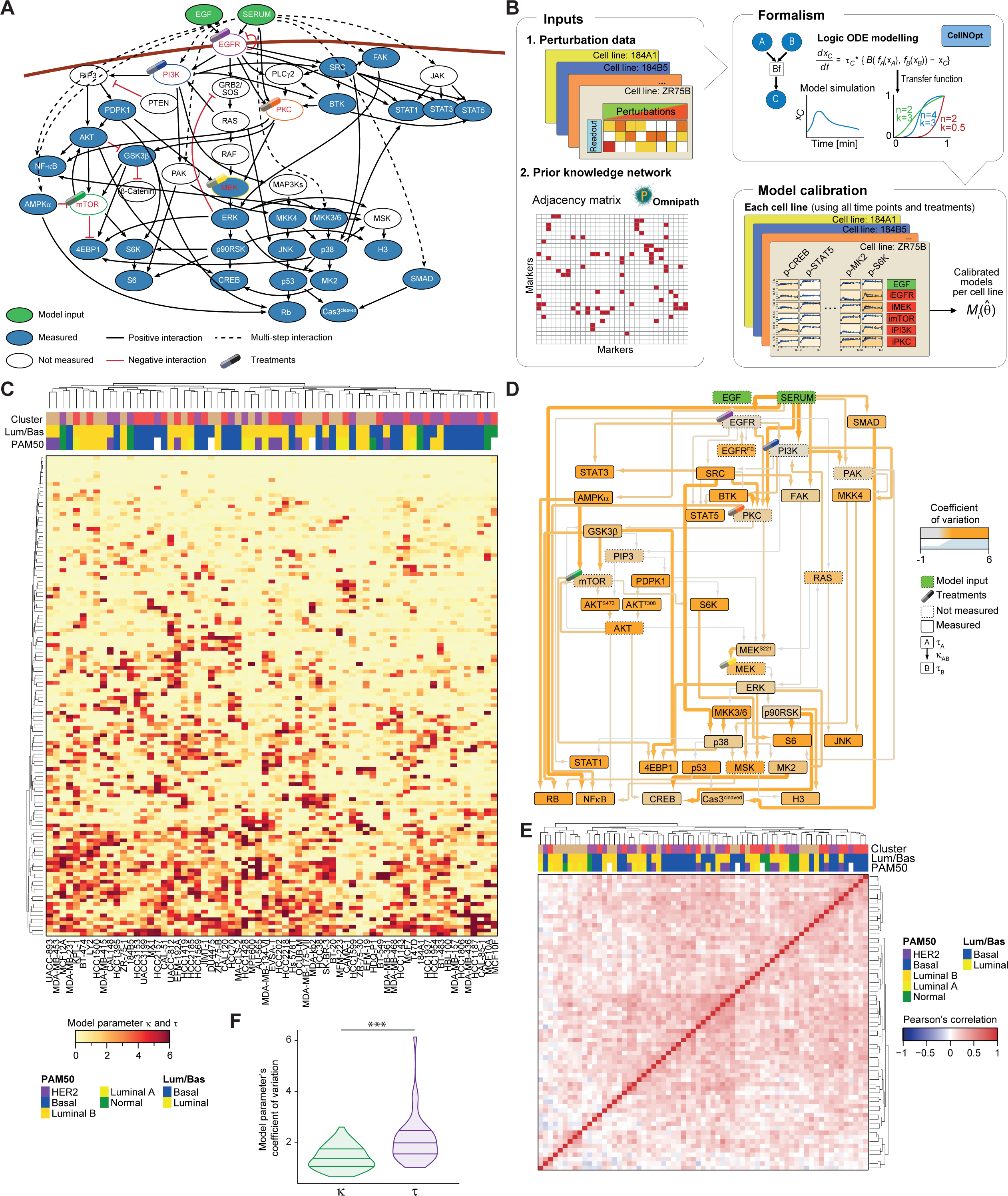
Cell-Line Specific Signaling Models. *(A) Prior knowledge network representation. Modeled but not measured nodes are indicated by white boxes, the model inputs are green, and intervention points are marked. Black, red, and dotted lines, respectively represent positive, negative, and multi-step interactions*. *(B) Computational approach for cell-line specific signaling model generation*. *(C) Clustering of the cell lines based on their model parameters τ (n = 40) and κ (n = 88). Color intensity on the white-yellow-red scale indicates raw parameter values. The PAM50 classification, luminal/basal classification, and information on the manually defined signaling cluster are shown. The data represents the model fitted on biological replicates*. *(D) The coefficient of variation values for κ and τ of the mechanistic signaling network models for all 67 cell lines are represented on the signaling network. The color and thickness of edges indicate κ parameter values on a low-to-high scale (gray-orange, thin-thick), and the node color indicates τ parameter values on a low-to-high scale (gray-orange). Modeled but not measured nodes are indicated by dotted boxes, the model inputs are green, and intervention points are marked*. *(E) Coefficient of variation across all cell lines plotted versus κ (green, n = 88) and τ (blue, n = 40). The horizontal lines represent the medians and the 25% and 75% quantiles (Welch two-sample t-test, two-sided, ***p ≤ 0.001)*. *(F) Clustering of the cell lines based on the Pearson’s correlation between κ and τ. Color intensity on a blue-white-red scale indicates Pearson’s correlation. The PAM50 classification, luminal/basal classification, and information on the manually defined signaling cluster are shown. The data represents the model fitted on biological replicates*.

**Supplementary Figure 4:**
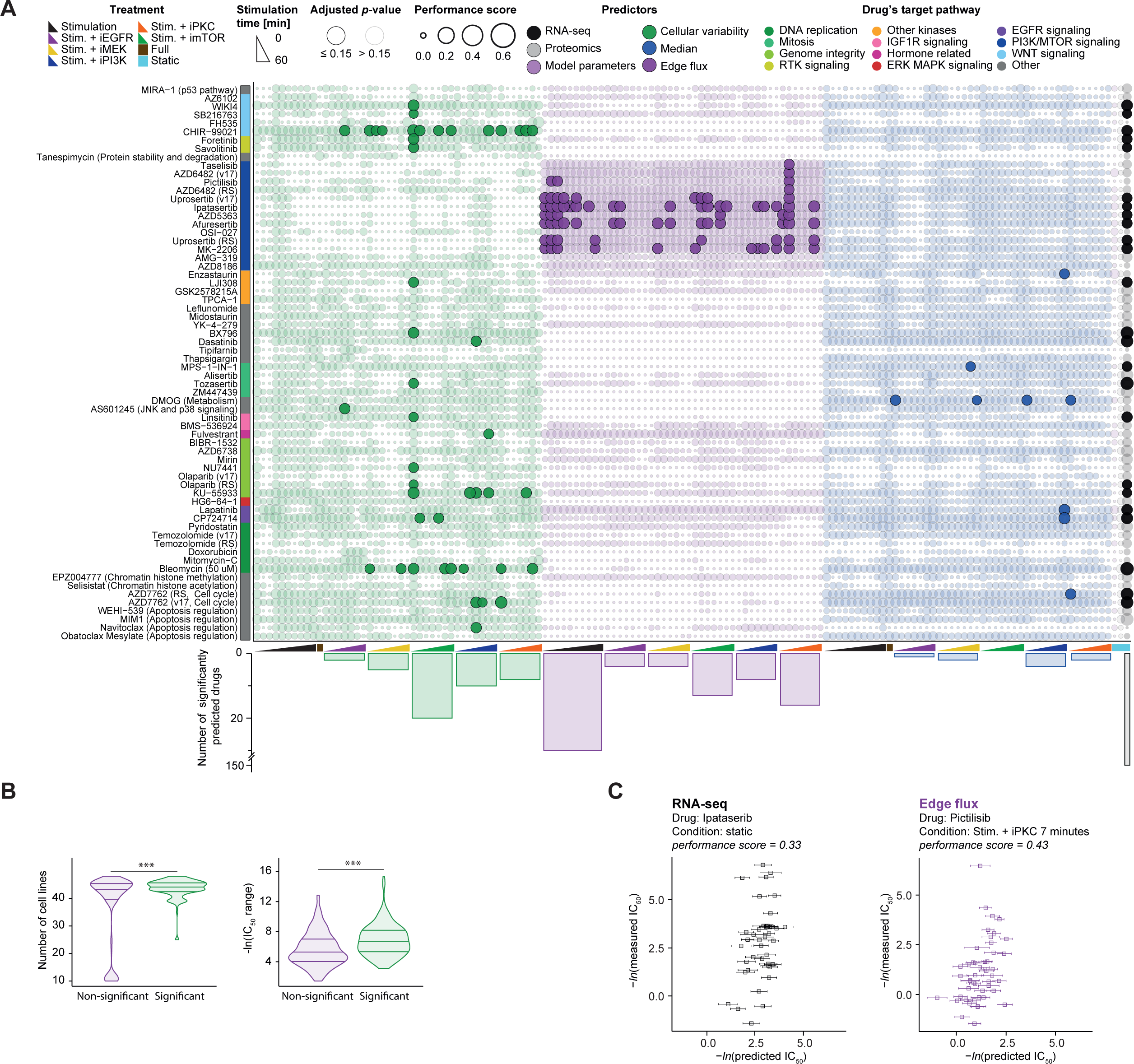
Prediction of Drug Sensitivity Using Dynamic Predictors. *(A) Upper: Drug sensitivities that are predicted by at least one dynamic predictor (FDR 25% and performance score > 0.2, multiple hypothesis correction for the 409 drug measurements) are shown in rows versus the predictors in columns, ordered by predictor, treatment, and increasing time. The bubble color indicates the predictor and size is proportional to the performance score. If the bubble circumference is black, the model for that predictor-drug combination was significant (FDR 15% and performance score > 0.3). Lower: Bar plot shows the number of drugs for which sensitivity was significantly and accurately predicted. The target pathway determines the drug arrangement (key at the top). Parentheses following the drug names give the version of the GDSC screen, if ambiguous*. *(B) Left: The number of cell lines for which drug predictions were non-significant (violet) or significantly accurate (green). Right: IC_50_ range in the training dataset for which drug predictions were non-significant (violet) or significantly accurate (green). The horizontal lines represent the medians and the 25% and 75% quantiles (n = 409, Welch two sample t-test, two-sided, ***p ≤ 0.001)*. *(C) Plots of measured and predicted IC_50_ values for two predictor, drug, and condition combinations. The error bars indicate the standard deviation from the 5-fold cross-validation*.

**Supplementary Figure 5:**
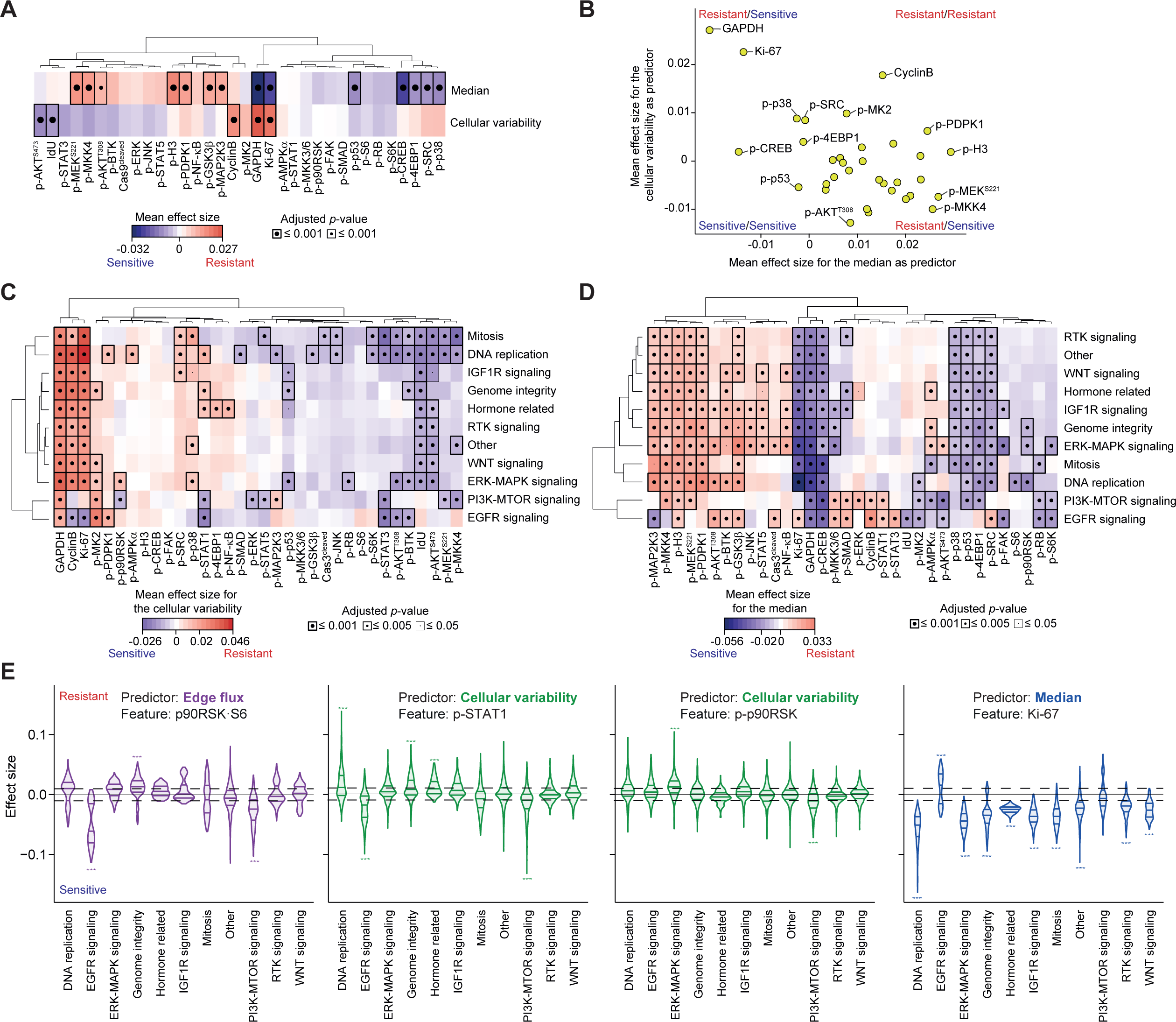
Features that Influence Drug Sensitivity Predictions. *(A) Mean effect size features of the cellular variability and median marker expression are shown. The features were hierarchically clustered. Mean effect sizes are indicated on a low-to-high color scale (blue-gray-red, sensitive-nonsignificant-resistant). The adjusted p-value is indicated by the dot size and the box thickness*. *(B) Mean effect size for the median marker expression as predictor plotted against the mean effect size for the cellular variability as predictor for matched features (n = 34). Interesting features are labeled*. *(C) Mean pathway-specific effect size features of the cellular variability as a predictor are shown. Drugs are binned according to the target pathway in rows, and features are shown in columns. Both the features and target pathways were hierarchically clustered. Mean effect sizes are indicated on a low-to-high color scale (blue-gray-red, sensitive-nonsignificant-resistant, FDR 5%). For each class the lowest adjusted p-value of all side-by-side comparisons is indicated by the dot size and the box thickness. The group “Other” contains all the drugs not falling into another group*. *(D) Mean pathway-specific effect size features of the median marker expression as a predictor are shown as described in panel C*. *(F) Effect size distributions for four selected features for which pathway-specific effects were observed. The selected predictors and features are indicated in each case. The dashed line indicates the significant threshold of 0.01 minimum effect size. The horizontal lines indicate the median and the 25% and 75% quantiles. Adjusted p-values, *p ≤ 0.05, **p ≤ 0.01, ***p ≤ 0.001*.

**Supplementary Figure 6:**
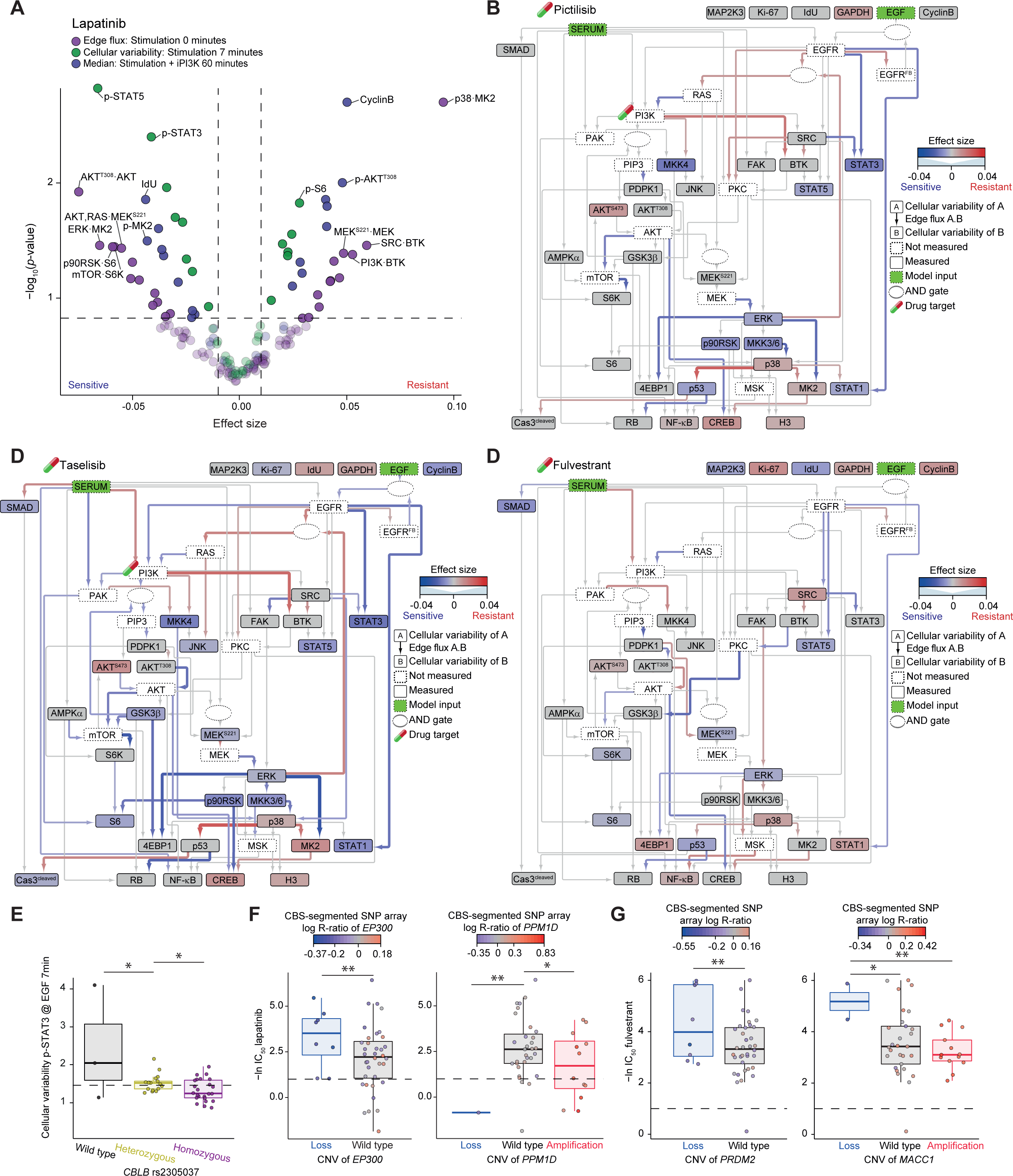
Resistance and Sensitivity to PI3K and EGFR Inhibition. *(A) Effect size vs. –log_10_ p-value for all features for the conditions with the highest performance score across cellular variability (stimulation 7 minutes), median (iPI3K 60 minutes), and edge flux (starvation). Non-significantly predicted features are drawn opaquely, the predictor is indicated by color, and the thresholds for significance as dashed lines (FDR 15% and effect size > 0.01 or < −0.01)*. *(B, C, D) Significant effect-size features of the cellular variability (nodes) and edge flux predictors (edges) represented on the signaling network for the drugs B) pictilisib, C) taselisib, and D) fulvestrant (FDR 15% and effect size > 0.01 or < −0.01). The effect sizes are indicated using color (nodes and edges), and the edge thicknesses are proportional to the absolute values of the effect sizes. An image of a drug capsule highlights the putative drug target. In each case the conditions with the highest performance scores are plotted: B, C) stimulation 9 minutes and PKC inhibitor 7 minutes for cellular variability and edge fluxes, respectively; and D) PI3K inhibitor 40 minutes and mTOR inhibitor 7 minutes for cellular variability and edge fluxes, respectively*. *(E) Values of the cellular variability of p-STAT3 (at stimulation 7 minutes) plotted against SNP status of rs2305037 (RNF213 gene, n = 61 cell lines). Thick lines indicate medians, the dashed line indicates the arbitrary threshold for high variability, boxes indicate the 25% and 75% quantiles, whiskers extend between the median and ± (1.58 * inter-quantile range). Each data point represents a cell line, and the color indicates the SNP status (ANOVA followed by Tukey honest significant differences computation: *p ≤ 0.3, **p ≤ 0.15, ***p ≤ 0.05)*. *(F) IC_50_ values (–ln) for lapatinib plotted against CNV status of EP300 or PPM1D (n = 42 cell lines). Plots and statistical analysis are as in panel E, except that the dashed line indicates the arbitrary threshold for sensitivity (1) and that color intensity indicates the amplification status on a low-to-high scale (blue-gray-red)*. *(G) IC_50_ values (–ln) for fulvestrant plotted against CNV status of PRDM2 or MACC1 (n = 42 cell lines). Plots and statistical analysis are as in panel E, except that the dashed line indicates the arbitrary threshold for sensitivity (1) and that color intensity indicates the amplification status on a low-to-high scale (blue-gray-red)*.

**Supplementary Table 1.**

| METAL TAG | TARGET | SUPPLIER | CLONE |
| --- | --- | --- | --- |
| **La139** | p-CREB | BD Biosciences | J151-21 |
| **Pr141** | p-STAT5 | Biolegend | 47/Stat5(Y694) |
| **Nd142** | p-SRC | eBioscience | SC1T2M3 |
| **Nd143** | p-FAK | Cell Signaling Technologies | polyclonal_Fak |
| **Nd144** | p-MEK1/2 | Cell Signaling Technologies | 166F8 |
| **Nd145** | p-MAPKAPK-2 | Cell Signaling Technologies | 27B7 |
| **Nd146** | p-S6K | Cell Signaling Technologies | 1A5 |
| **Sm147** | p-MAP2K3 | Assay Biotech | polyclonal_16 |
| **Nd148** | p-STAT1 | Cell Signaling Technologies | Ser727 |
| **Sm149** | p-p53 | Cell Signaling Technologies | 16G8 |
| **Nd150** | p-NF-κB | Biolegend | K10-895.12.50 |
| **Eu151** | p-p38 | BD Biosciences | 36/p38<br>(pT180/pY182) |
| **Sm152** | p-AMPKα | Cell Signaling Technologies | 40H9 |
| **Eu153** | p-AKT(473) | Cell Signaling Technologies | D9E |
| **Sm154** | p-ERK1/2 | BD Biosciences | 20A |
| **Gd156** | Cyclin B1 | BD Biosciences | GNS-11 |
| **Gd158** | p-GSK3β | Cell Signaling Technologies | D85E12 |
| **Tb159** | GAPDH | Thermo Scientific Pierce Antibodies | 6C5 |
| **Gd160** | p-MKK3-MKK6 | Cell Signaling Technologies | D8E9 |
| **Dy161** | p-PDPK1 | BD Biosciences | J66-653.44.22 |
| **Dy162** | p-BTK | BD Biosciences | 24a/BTK |
| **Dy163** | p-p90RSK | Cell Signaling Technologies | D5D8 |
| **Dy164** | p-SMAD2/3 | Cell Signaling Technologies | D27F4 |
| **Ho165** | β-CATENIN | Cell Signaling Technologies | D13A1 |
| **Er166** | p-STAT3 | BD Biosciences | 4/pStat3 |
| **Er167** | p-SAPK/JNK | Cell Signaling Technologies | G9 |
| **Er168** | Ki-67 | BD Biosciences | B56 |
| **Er170** | p-Histone H3 | Biolegend | HTA28 |
| **Yb171** | p-S6 | BD Biosciences | N7-548 |
| **Yb172** | cleaved PARP | BD Biosciences | F21-852 |
| **Yb172** | Cleaved Caspase3 | BD Biosciences | C92-605 |
| **Yb173** | p-SEK/MKK4 | Cell Signaling Technologies | C36C11 |
| **Yb174** | p-AKT(308) | Cell Signaling Technologies | D25E6 |
| **Lu175** | p-RB | Cell Signaling Technologies | D20B12 |
| **Yb176** | p-4EBP1 | Cell Signaling Technologies | 236B4 |

**Supplementary Table 2.**

| TREATMENT | REPLICATE A [MIN] | REPLICATE B [MIN] |
| --- | --- | --- |
| Stimulation with EGF and FBS | 0, 5.5, 9, 13, 23, 40 | 0, 7, 9, 17, 30, 60 |
| Stimulation with EGF and FBS + iPI3K | 0, 9, 13, 40 | 0, 7, 17, 60 |
| Stimulation with EGF and FBS + imTOR | 0, 9, 13, 40 | 0, 7, 17, 60 |
| Stimulation with EGF and FBS + iEGFR | 0, 9, 13, 40 | 0, 7, 17, 60 |
| Stimulation with EGF and FBS + iPKC | 0, 9, 13, 40 | 0, 7, 17, 60 |
| Stimulation with EGF and FBS + iMEK | 0, 9, 13, 40 | 0, 7, 17, 60 |
| No stimulation | 0 | 0 |
| Biological control (cell line HCC70) | 0, 15 | 0, 10 |

**Supplementary Table 3.**

| TARGET MOLECULE | INHIBITOR NAME | ALTERNATIVE NAME | SUPPLIER | CATALOG NUMBER | FINAL CONCENTRATION USED IN μM |
| --- | --- | --- | --- | --- | --- |
| PI3K | GDC-0941 | Pictilisib | LC Laboratories | G-9252 | 0.5 |
| mTOR | Rapamycin | Sirolimus | LC Laboratories | R-5000 | 0.01 |
| EGFR | Lapatinib |  | LC Laboratories | L-4899 | 1.08 |
| PKC | Enzastaurin | LY317615 | LC Laboratories | E-4506 | 3.9 |
| MEK | CI-1040 | PD184352 | Selleckchem | S1020 | 1.7 |

**Supplementary Table 4.**
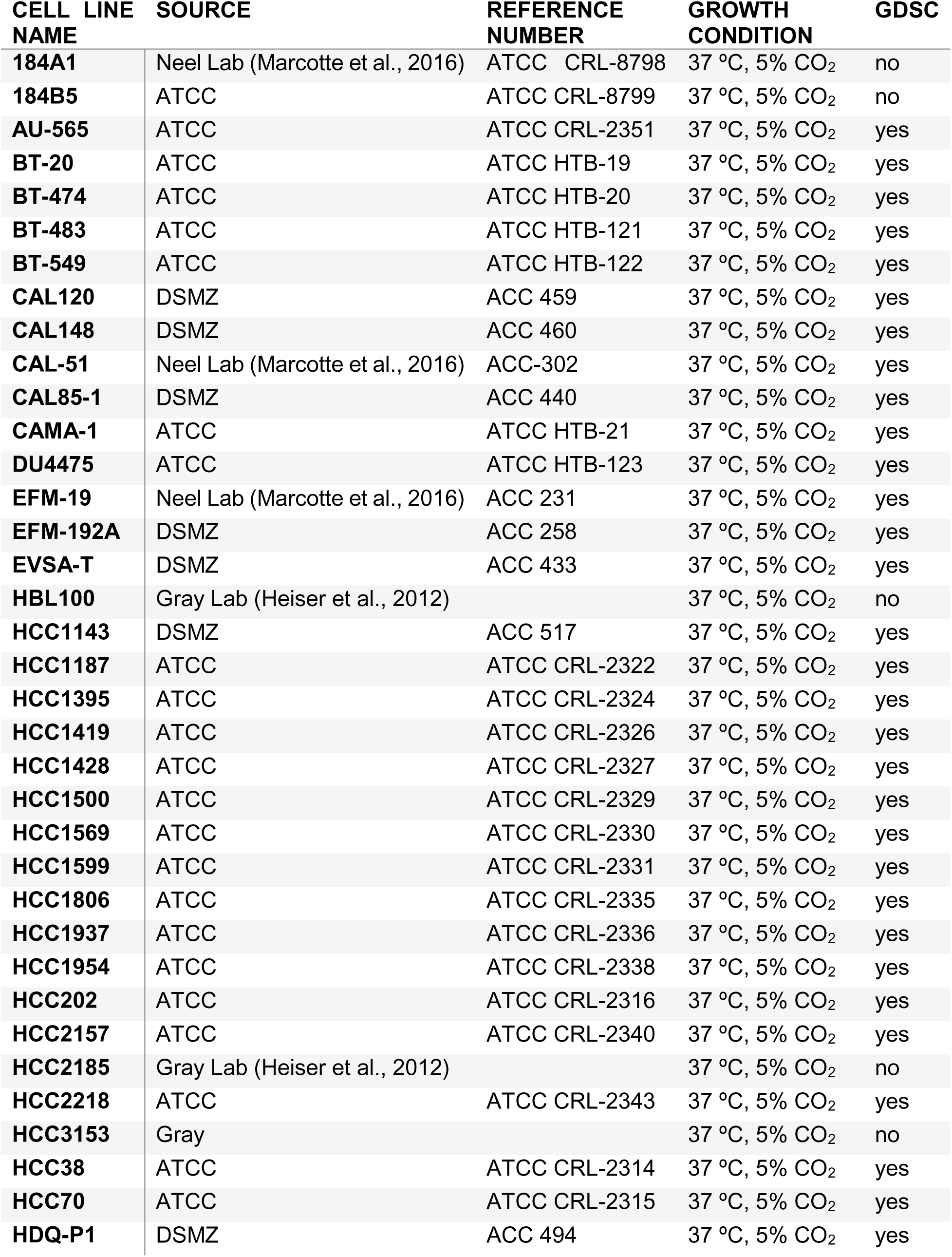
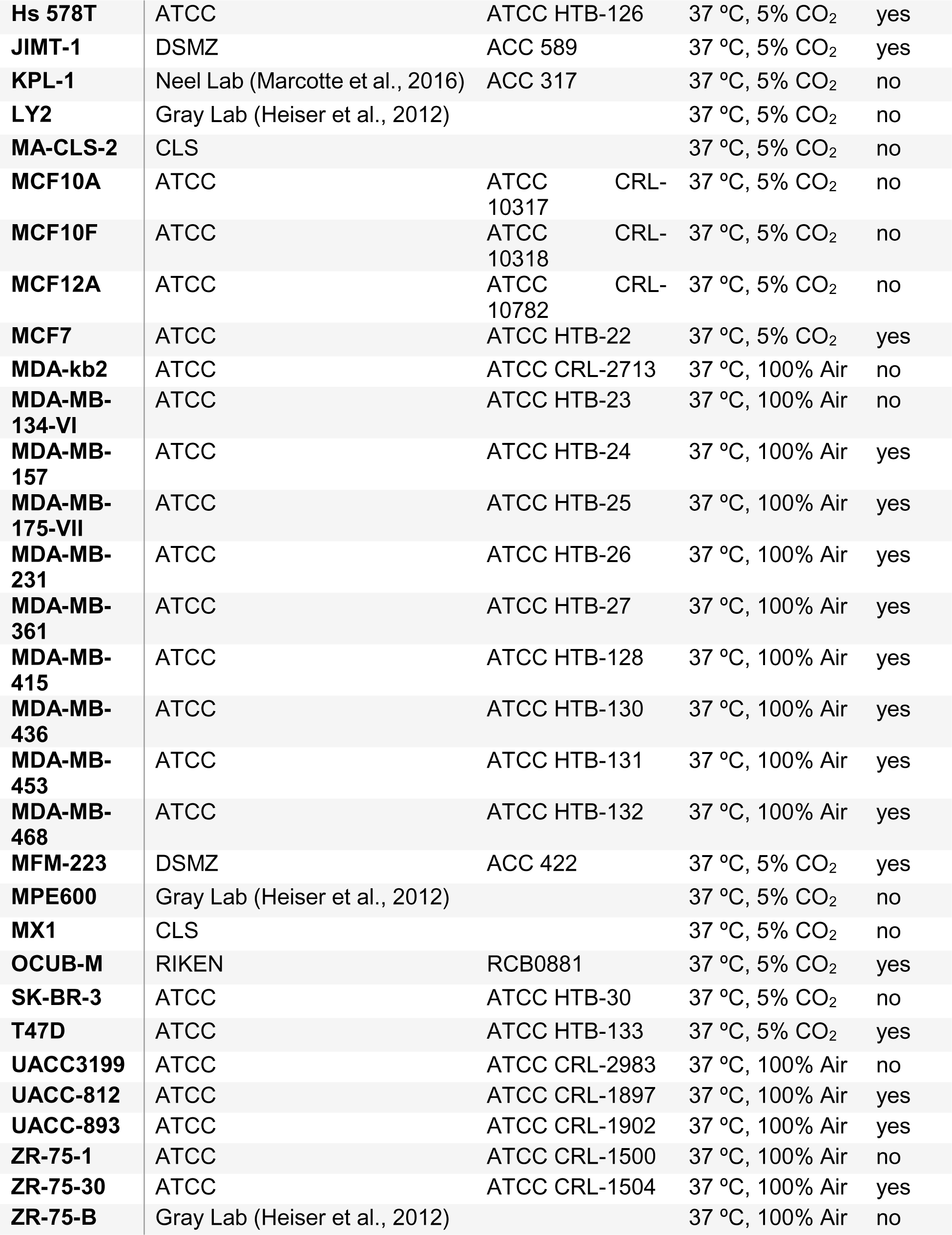

**Supplementary Table 5.**
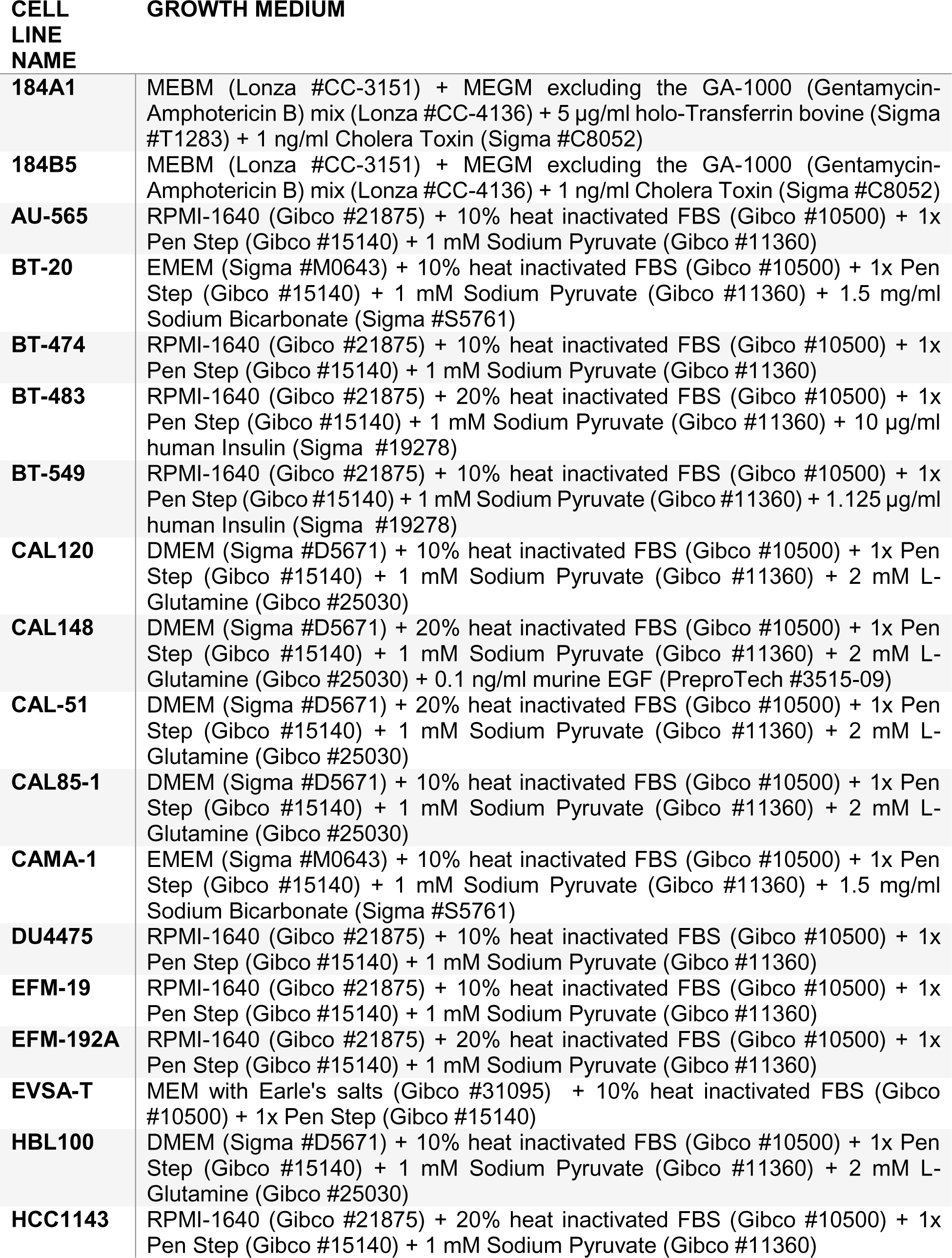
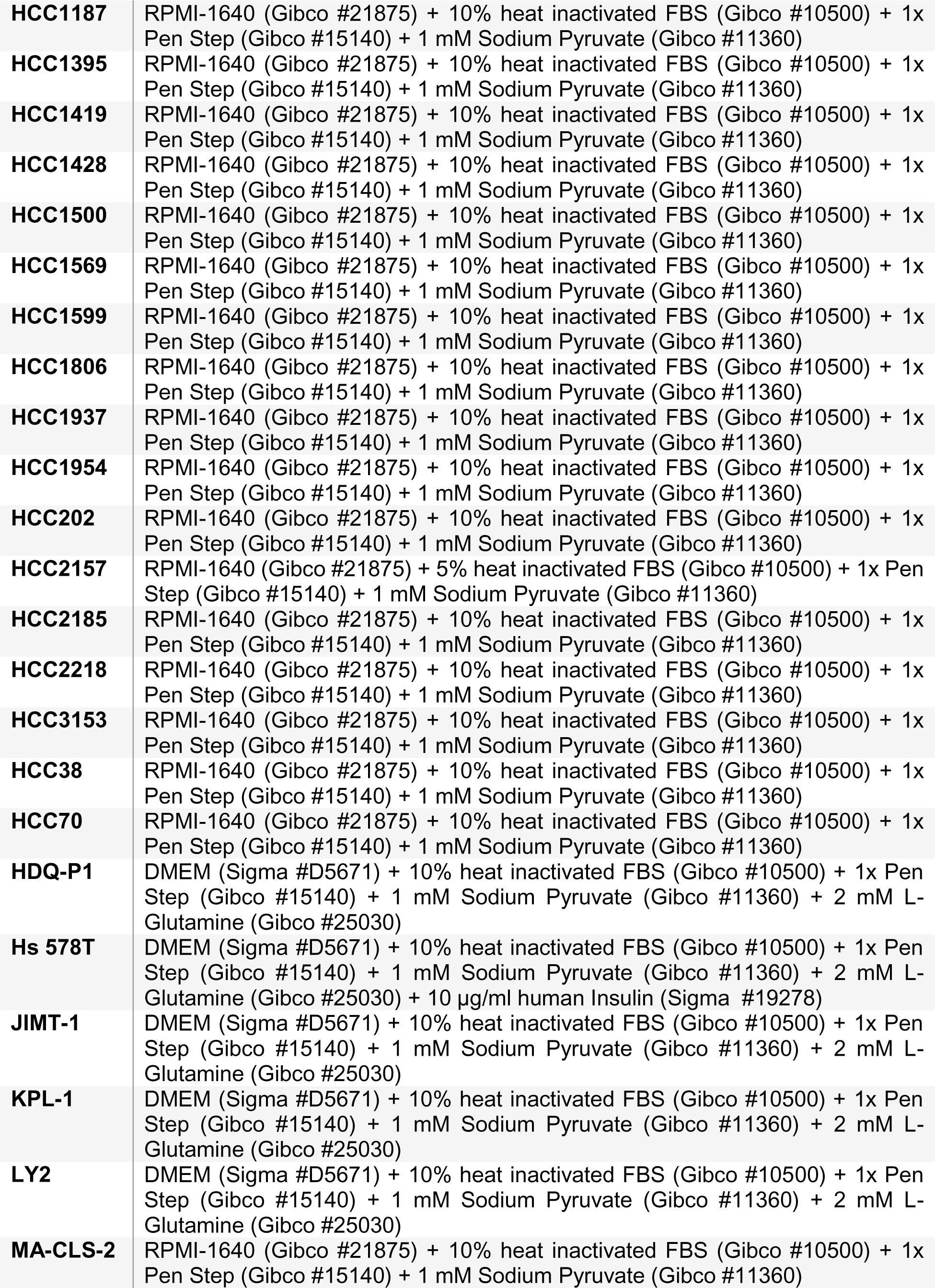
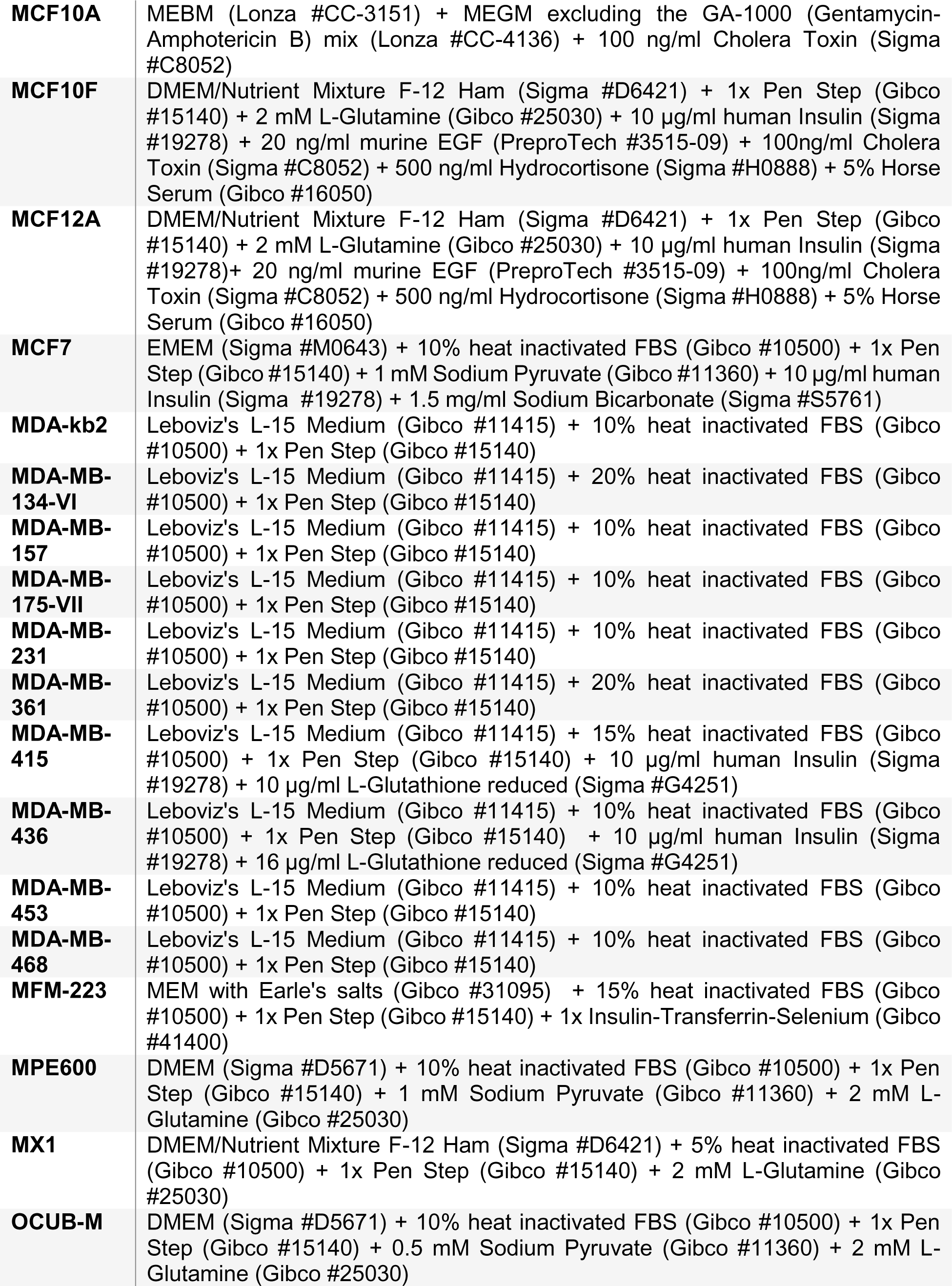
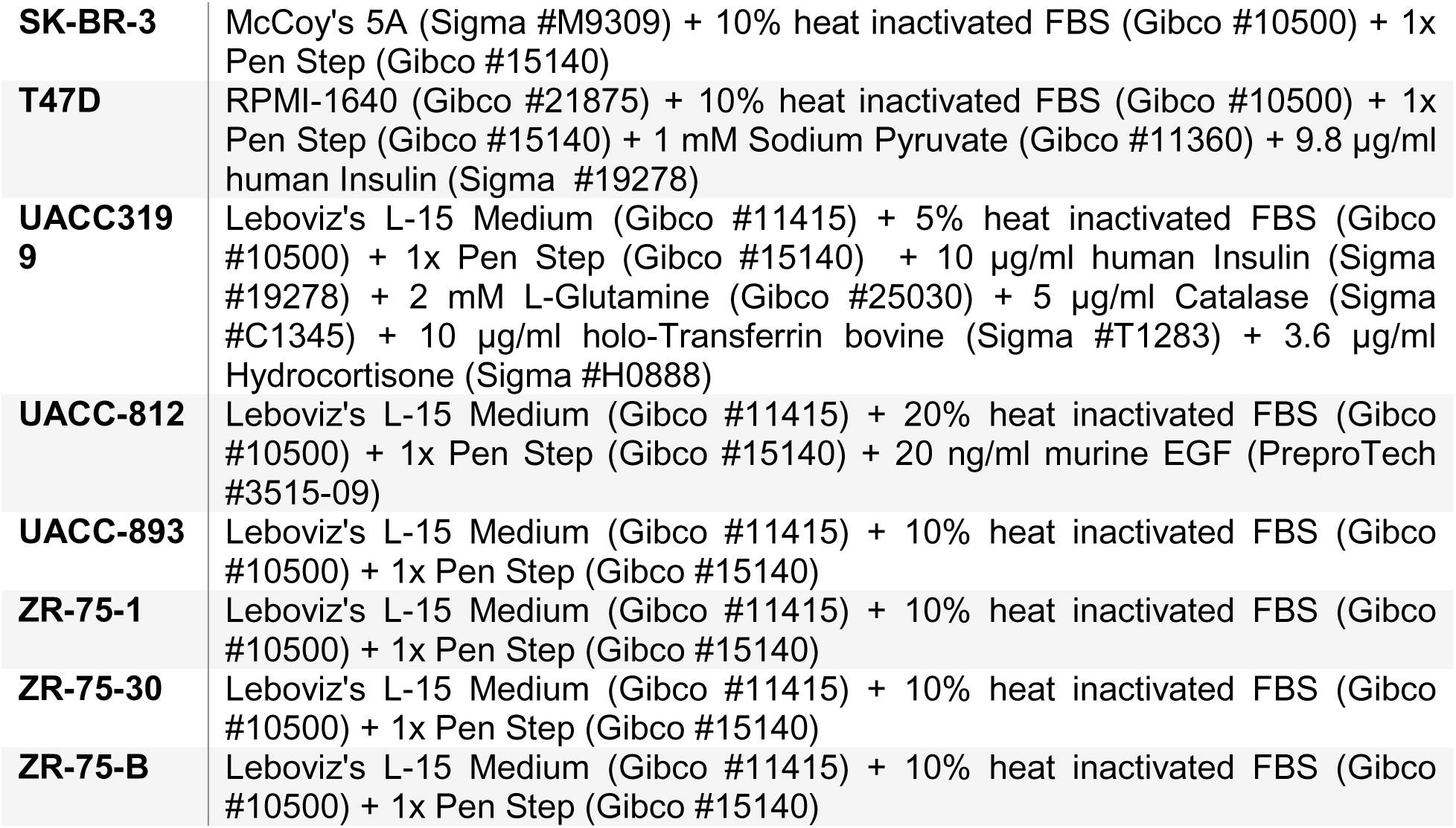

| CELL LINE NAME | GROWTH MEDIUM |
| --- | --- |
| **184A1** | MEBM (Lonza #CC-3151) + MEGM excluding the GA-1000 (Gentamycin-Amphotericin B) mix (Lonza #CC-4136) + 5 µg/ml holo-Transferrin bovine (Sigma #T1283) + 1 ng/ml Cholera Toxin (Sigma #C8052) |
| **184B5** | MEBM (Lonza #CC-3151) + MEGM excluding the GA-1000 (Gentamycin-Amphotericin B) mix (Lonza #CC-4136) + 1 ng/ml Cholera Toxin (Sigma #C8052) |
| **AU-565** | RPMI-1640 (Gibco #21875) + 10% heat inactivated FBS (Gibco #10500) + 1x Pen Step (Gibco #15140) + 1 mM Sodium Pyruvate (Gibco #11360) |
| **BT-20** | EMEM (Sigma #M0643) + 10% heat inactivated FBS (Gibco #10500) + 1x Pen Step (Gibco #15140) + 1 mM Sodium Pyruvate (Gibco #11360) + 1.5 mg/ml Sodium Bicarbonate (Sigma #S5761) |
| **BT-474** | RPMI-1640 (Gibco #21875) + 10% heat inactivated FBS (Gibco #10500) + 1x Pen Step (Gibco #15140) + 1 mM Sodium Pyruvate (Gibco #11360) |
| **BT-483** | RPMI-1640 (Gibco #21875) + 20% heat inactivated FBS (Gibco #10500) + 1x Pen Step (Gibco #15140) + 1 mM Sodium Pyruvate (Gibco #11360) + 10 µg/ml human Insulin (Sigma #19278) |
| **BT-549** | RPMI-1640 (Gibco #21875) + 10% heat inactivated FBS (Gibco #10500) + 1x Pen Step (Gibco #15140) + 1 mM Sodium Pyruvate (Gibco #11360) + 1.125 µg/ml human Insulin (Sigma #19278) |
| **CAL120** | DMEM (Sigma #D5671) + 10% heat inactivated FBS (Gibco #10500) + 1x Pen Step (Gibco #15140) + 1 mM Sodium Pyruvate (Gibco #11360) + 2 mM L-Glutamine (Gibco #25030) |
| **CAL148** | DMEM (Sigma #D5671) + 20% heat inactivated FBS (Gibco #10500) + 1x Pen Step (Gibco #15140) + 1 mM Sodium Pyruvate (Gibco #11360) + 2 mM L-Glutamine (Gibco #25030) + 0.1 ng/ml murine EGF (PreproTech #3515-09) |
| **CAL-51** | DMEM (Sigma #D5671) + 20% heat inactivated FBS (Gibco #10500) + 1x Pen Step (Gibco #15140) + 1 mM Sodium Pyruvate (Gibco #11360) + 2 mM L-Glutamine (Gibco #25030) |
| **CAL85-1** | DMEM (Sigma #D5671) + 10% heat inactivated FBS (Gibco #10500) + 1x Pen Step (Gibco #15140) + 1 mM Sodium Pyruvate (Gibco #11360) + 2 mM L-Glutamine (Gibco #25030) |
| **CAMA-1** | EMEM (Sigma #M0643) + 10% heat inactivated FBS (Gibco #10500) + 1x Pen Step (Gibco #15140) + 1 mM Sodium Pyruvate (Gibco #11360) + 1.5 mg/ml Sodium Bicarbonate (Sigma #S5761) |
| **DU4475** | RPMI-1640 (Gibco #21875) + 10% heat inactivated FBS (Gibco #10500) + 1x Pen Step (Gibco #15140) + 1 mM Sodium Pyruvate (Gibco #11360) |
| **EFM-19** | RPMI-1640 (Gibco #21875) + 10% heat inactivated FBS (Gibco #10500) + 1x Pen Step (Gibco #15140) + 1 mM Sodium Pyruvate (Gibco #11360) |
| **EFM-192A** | RPMI-1640 (Gibco #21875) + 20% heat inactivated FBS (Gibco #10500) + 1x Pen Step (Gibco #15140) + 1 mM Sodium Pyruvate (Gibco #11360) |
| **EVSA-T** | MEM with Earle's salts (Gibco #31095) + 10% heat inactivated FBS (Gibco #10500) + 1x Pen Step (Gibco #15140) |
| **HBL100** | DMEM (Sigma #D5671) + 10% heat inactivated FBS (Gibco #10500) + 1x Pen Step (Gibco #15140) + 1 mM Sodium Pyruvate (Gibco #11360) + 2 mM L-Glutamine (Gibco #25030) |
| **HCC1143** | RPMI-1640 (Gibco #21875) + 20% heat inactivated FBS (Gibco #10500) + 1x Pen Step (Gibco #15140) + 1 mM Sodium Pyruvate (Gibco #11360) |
| **HCC1187** | RPMI-1640 (Gibco #21875) + 10% heat inactivated FBS (Gibco #10500) + 1x Pen Step (Gibco #15140) + 1 mM Sodium Pyruvate (Gibco #11360) |
| **HCC1395** | RPMI-1640 (Gibco #21875) + 10% heat inactivated FBS (Gibco #10500) + 1x Pen Step (Gibco #15140) + 1 mM Sodium Pyruvate (Gibco #11360) |
| **HCC1419** | RPMI-1640 (Gibco #21875) + 10% heat inactivated FBS (Gibco #10500) + 1x Pen Step (Gibco #15140) + 1 mM Sodium Pyruvate (Gibco #11360) |
| **HCC1428** | RPMI-1640 (Gibco #21875) + 10% heat inactivated FBS (Gibco #10500) + 1x Pen Step (Gibco #15140) + 1 mM Sodium Pyruvate (Gibco #11360) |
| **HCC1500** | RPMI-1640 (Gibco #21875) + 10% heat inactivated FBS (Gibco #10500) + 1x Pen Step (Gibco #15140) + 1 mM Sodium Pyruvate (Gibco #11360) |
| **HCC1569** | RPMI-1640 (Gibco #21875) + 10% heat inactivated FBS (Gibco #10500) + 1x Pen Step (Gibco #15140) + 1 mM Sodium Pyruvate (Gibco #11360) |
| **HCC1599** | RPMI-1640 (Gibco #21875) + 10% heat inactivated FBS (Gibco #10500) + 1x Pen Step (Gibco #15140) + 1 mM Sodium Pyruvate (Gibco #11360) |
| **HCC1806** | RPMI-1640 (Gibco #21875) + 10% heat inactivated FBS (Gibco #10500) + 1x Pen Step (Gibco #15140) + 1 mM Sodium Pyruvate (Gibco #11360) |
| **HCC1937** | RPMI-1640 (Gibco #21875) + 10% heat inactivated FBS (Gibco #10500) + 1x Pen Step (Gibco #15140) + 1 mM Sodium Pyruvate (Gibco #11360) |
| **HCC1954** | RPMI-1640 (Gibco #21875) + 10% heat inactivated FBS (Gibco #10500) + 1x Pen Step (Gibco #15140) + 1 mM Sodium Pyruvate (Gibco #11360) |
| **HCC202** | RPMI-1640 (Gibco #21875) + 10% heat inactivated FBS (Gibco #10500) + 1x Pen Step (Gibco #15140) + 1 mM Sodium Pyruvate (Gibco #11360) |
| **HCC2157** | RPMI-1640 (Gibco #21875) + 5% heat inactivated FBS (Gibco #10500) + 1x Pen Step (Gibco #15140) + 1 mM Sodium Pyruvate (Gibco #11360) |
| **HCC2185** | RPMI-1640 (Gibco #21875) + 10% heat inactivated FBS (Gibco #10500) + 1x Pen Step (Gibco #15140) + 1 mM Sodium Pyruvate (Gibco #11360) |
| **HCC2218** | RPMI-1640 (Gibco #21875) + 10% heat inactivated FBS (Gibco #10500) + 1x Pen Step (Gibco #15140) + 1 mM Sodium Pyruvate (Gibco #11360) |
| **HCC3153** | RPMI-1640 (Gibco #21875) + 10% heat inactivated FBS (Gibco #10500) + 1x Pen Step (Gibco #15140) + 1 mM Sodium Pyruvate (Gibco #11360) |
| **HCC38** | RPMI-1640 (Gibco #21875) + 10% heat inactivated FBS (Gibco #10500) + 1x Pen Step (Gibco #15140) + 1 mM Sodium Pyruvate (Gibco #11360) |
| **HCC70** | RPMI-1640 (Gibco #21875) + 10% heat inactivated FBS (Gibco #10500) + 1x Pen Step (Gibco #15140) + 1 mM Sodium Pyruvate (Gibco #11360) |
| **HDQ-P1** | DMEM (Sigma #D5671) + 10% heat inactivated FBS (Gibco #10500) + 1x Pen Step (Gibco #15140) + 1 mM Sodium Pyruvate (Gibco #11360) + 2 mM L-Glutamine (Gibco #25030) |
| **Hs 578T** | DMEM (Sigma #D5671) + 10% heat inactivated FBS (Gibco #10500) + 1x Pen Step (Gibco #15140) + 1 mM Sodium Pyruvate (Gibco #11360) + 2 mM L-Glutamine (Gibco #25030) + 10 µg/ml human Insulin (Sigma #19278) |
| **JIMT-1** | DMEM (Sigma #D5671) + 10% heat inactivated FBS (Gibco #10500) + 1x Pen Step (Gibco #15140) + 1 mM Sodium Pyruvate (Gibco #11360) + 2 mM L-Glutamine (Gibco #25030) |
| **KPL-1** | DMEM (Sigma #D5671) + 10% heat inactivated FBS (Gibco #10500) + 1x Pen Step (Gibco #15140) + 1 mM Sodium Pyruvate (Gibco #11360) + 2 mM L-Glutamine (Gibco #25030) |
| **LY2** | DMEM (Sigma #D5671) + 10% heat inactivated FBS (Gibco #10500) + 1x Pen Step (Gibco #15140) + 1 mM Sodium Pyruvate (Gibco #11360) + 2 mM L-Glutamine (Gibco #25030) |
| **MA-CLS-2** | RPMI-1640 (Gibco #21875) + 10% heat inactivated FBS (Gibco #10500) + 1x Pen Step (Gibco #15140) + 1 mM Sodium Pyruvate (Gibco #11360) |
| **MCF10A** | MEBM (Lonza #CC-3151) + MEGM excluding the GA-1000 (Gentamycin-Amphotericin B) mix (Lonza #CC-4136) + 100 ng/ml Cholera Toxin (Sigma #C8052) |
| **MCF10F** | DMEM/Nutrient Mixture F-12 Ham (Sigma #D6421) + 1x Pen Step (Gibco #15140) + 2 mM L-Glutamine (Gibco #25030) + 10 µg/ml human Insulin (Sigma #19278) + 20 ng/ml murine EGF (PreproTech #3515-09) + 100ng/ml Cholera Toxin (Sigma #C8052) + 500 ng/ml Hydrocortisone (Sigma #H0888) + 5% Horse Serum (Gibco #16050) |
| **MCF12A** | DMEM/Nutrient Mixture F-12 Ham (Sigma #D6421) + 1x Pen Step (Gibco #15140) + 2 mM L-Glutamine (Gibco #25030) + 10 µg/ml human Insulin (Sigma #19278)+ 20 ng/ml murine EGF (PreproTech #3515-09) + 100ng/ml Cholera Toxin (Sigma #C8052) + 500 ng/ml Hydrocortisone (Sigma #H0888) + 5% Horse Serum (Gibco #16050) |
| **MCF7** | EMEM (Sigma #M0643) + 10% heat inactivated FBS (Gibco #10500) + 1x Pen Step (Gibco #15140) + 1 mM Sodium Pyruvate (Gibco #11360) + 10 µg/ml human Insulin (Sigma #19278) + 1.5 mg/ml Sodium Bicarbonate (Sigma #S5761) |
| **MDA-kb2** | Leboviz's L-15 Medium (Gibco #11415) + 10% heat inactivated FBS (Gibco #10500) + 1x Pen Step (Gibco #15140) |
| **MDA-MB-134-VI** | Leboviz's L-15 Medium (Gibco #11415) + 20% heat inactivated FBS (Gibco #10500) + 1x Pen Step (Gibco #15140) |
| **MDA-MB-157** | Leboviz's L-15 Medium (Gibco #11415) + 10% heat inactivated FBS (Gibco #10500) + 1x Pen Step (Gibco #15140) |
| **MDA-MB-175-VII** | Leboviz's L-15 Medium (Gibco #11415) + 10% heat inactivated FBS (Gibco #10500) + 1x Pen Step (Gibco #15140) |
| **MDA-MB-231** | Leboviz's L-15 Medium (Gibco #11415) + 10% heat inactivated FBS (Gibco #10500) + 1x Pen Step (Gibco #15140) |
| **MDA-MB-361** | Leboviz's L-15 Medium (Gibco #11415) + 20% heat inactivated FBS (Gibco #10500) + 1x Pen Step (Gibco #15140) |
| **MDA-MB-415** | Leboviz's L-15 Medium (Gibco #11415) + 15% heat inactivated FBS (Gibco #10500) + 1x Pen Step (Gibco #15140) + 10 µg/ml human Insulin (Sigma #19278) + 10 µg/ml L-Glutathione reduced (Sigma #G4251) |
| **MDA-MB-436** | Leboviz's L-15 Medium (Gibco #11415) + 10% heat inactivated FBS (Gibco #10500) + 1x Pen Step (Gibco #15140) + 10 µg/ml human Insulin (Sigma #19278) + 16 µg/ml L-Glutathione reduced (Sigma #G4251) |
| **MDA-MB-453** | Leboviz's L-15 Medium (Gibco #11415) + 10% heat inactivated FBS (Gibco #10500) + 1x Pen Step (Gibco #15140) |
| **MDA-MB-468** | Leboviz's L-15 Medium (Gibco #11415) + 10% heat inactivated FBS (Gibco #10500) + 1x Pen Step (Gibco #15140) |
| **MFM-223** | MEM with Earle's salts (Gibco #31095) + 15% heat inactivated FBS (Gibco #10500) + 1x Pen Step (Gibco #15140) + 1x Insulin-Transferrin-Selenium (Gibco #41400) |
| **MPE600** | DMEM (Sigma #D5671) + 10% heat inactivated FBS (Gibco #10500) + 1x Pen Step (Gibco #15140) + 1 mM Sodium Pyruvate (Gibco #11360) + 2 mM L-Glutamine (Gibco #25030) |
| **MX1** | DMEM/Nutrient Mixture F-12 Ham (Sigma #D6421) + 5% heat inactivated FBS (Gibco #10500) + 1x Pen Step (Gibco #15140) + 2 mM L-Glutamine (Gibco #25030) |
| **OCUB-M** | DMEM (Sigma #D5671) + 10% heat inactivated FBS (Gibco #10500) + 1x Pen Step (Gibco #15140) + 0.5 mM Sodium Pyruvate (Gibco #11360) + 2 mM L-Glutamine (Gibco #25030) |
| **SK-BR-3** | McCoy's 5A (Sigma #M9309) + 10% heat inactivated FBS (Gibco #10500) + 1x Pen Step (Gibco #15140) |
| **T47D** | RPMI-1640 (Gibco #21875) + 10% heat inactivated FBS (Gibco #10500) + 1x Pen Step (Gibco #15140) + 1 mM Sodium Pyruvate (Gibco #11360) + 9.8 µg/ml human Insulin (Sigma #19278) |
| **UACC319**<br>**9** | Leboviz's L-15 Medium (Gibco #11415) + 5% heat inactivated FBS (Gibco #10500) + 1x Pen Step (Gibco #15140) + 10 µg/ml human Insulin (Sigma #19278) + 2 mM L-Glutamine (Gibco #25030) + 5 µg/ml Catalase (Sigma #C1345) + 10 µg/ml holo-Transferrin bovine (Sigma #T1283) + 3.6 µg/ml Hydrocortisone (Sigma #H0888) |
| **UACC-812** | Leboviz's L-15 Medium (Gibco #11415) + 20% heat inactivated FBS (Gibco #10500) + 1x Pen Step (Gibco #15140) + 20 ng/ml murine EGF (PreproTech #3515-09) |
| **UACC-893** | Leboviz's L-15 Medium (Gibco #11415) + 10% heat inactivated FBS (Gibco #10500) + 1x Pen Step (Gibco #15140) |
| **ZR-75-1** | Leboviz's L-15 Medium (Gibco #11415) + 10% heat inactivated FBS (Gibco #10500) + 1x Pen Step (Gibco #15140) |
| **ZR-75-30** | Leboviz's L-15 Medium (Gibco #11415) + 10% heat inactivated FBS (Gibco #10500) + 1x Pen Step (Gibco #15140) |
| **ZR-75-B** | Leboviz's L-15 Medium (Gibco #11415) + 10% heat inactivated FBS (Gibco #10500) + 1x Pen Step (Gibco #15140) |

**Supplementary Table 6.**

| SOURCE<br>NODE | INTERACTION<br>TYPE | TARGET<br>NODE |
| --- | --- | --- |
| PIP3 | 1 | AKT<sup>S473</sup> |
| p53 | 1 | RB |
| GSK3β | 1 | RB |
| AMPKα | 1 | RB |
| p53 | 1 | Cas3<sup>cleaved</sup> |
| SMAD | 1 | Cas3<sup>cleaved</sup> |
| ERK | 1 | MSK |
| MKK3/6 | 1 | MSK |
| MSK | 1 | H3 |
| p90RSK | 1 | H3 |
| SERUM | 1 | SMAD |
| SERUM | 1 | AMPKα |
| AMPKα | -1 | mTOR |
| AKT | 1 | NF-κB |
| SERUM | 1 | cAMP |
| cAMP | 1 | PKA |
| PKA | 1 | NF-κB |
| PKC | 1 | NF-κB |
| p38 | 1 | MSK |
| MSK | 1 | NF-κB |
| PLCγ2 | 1 | PKC |
| MET | 1 | PLCγ2 |
| PKC | 1 | RAF |
| PKC | 1 | GSK3β |
| mTOR | 1 | S6K |
| mTOR | 1 | AKT<sup>S473</sup> |
| PDPK1 | 1 | AKT<sup>T308</sup> |
| AKT<sup>S473</sup> | 1 | AKT |
| AKT<sup>T308</sup> | 1 | AKT |
| p38 | 1 | p53 |
| p38 | 1 | MK2 |
| GSK3β | 1 | PTEN |
| GSK3β | 1 | mTOR |
| GSK3β | 1 | β-Catenin |
| S6K | 1 | S6 |
| p90RSK | 1 | CREB |
| p90RSK | 1 | S6 |
| ERK | 1 | MKK3/6 |
| ERK | 1 | MK2 |
| ERK | 1 | ERK<sup>dm2</sup> |
| RAS | 1 | PI3K |
| BTK | 1 | PLCγ2 |
| AKT | 1 | mTOR |
| AKT | 1 | GSK3β |
| AKT | 1 | RAF<sup>S259</sup> |
| AKT | 1 | CREB |
| MK2 | 1 | CREB |
| PIP3 | 1 | PDPK1 |
| PDPK1 | 1 | S6K |
| PKC | 1 | MARCKS |
| SRC | 1 | PLCγ2 |
| SRC | 1 | BTK |
| SRC | 1 | FAK |
| EGFR | 1 | PI3K |
| EGFR | 1 | PLCγ2 |
| EGFR | 1 | EGFR<sup>FB</sup> |
| PDPK1 | 1 | MEK<sup>S221</sup> |
| RAF | 1 | MEK<sup>S221</sup> |
| mTOR | 1 | 4EBP1 |
| GSK3β | 1 | 4EBP1 |
| ERK | 1 | 4EBP1 |
| p38 | 1 | 4EBP1 |
| ERK | 1 | p90RSK |
| PI3K | 1 | SYK |
| SYK | 1 | BTK |
| MKK4 | 1 | JNK |
| PI3K | 1 | MAP3Ks |
| INSR | 1 | PI3K |
| PI3K | 1 | PAK |
| PAK | 1 | S6 |
| MET | 1 | PI3K |
| MET | 1 | FAK |
| BTK | 1 | STAT5 |
| EGFR | 1 | and1 |
| ERK<sup>dm2</sup> | -1 | and1 |
| and1 | 1 | RAS |
| PTEN | -1 | and2 |
| PI3K | 1 | and2 |
| and2 | 1 | PIP3 |
| EGFR<sup>FB</sup> | -1 | and3 |
| EGF | 1 | and3 |
| and3 | 1 | EGFR |
| RAS | 1 | and4 |
| RAF<sup>S259</sup> | -1 | and4 |
| and4 | 1 | RAF |
| MEK<sup>S221</sup> | 1 | MEK |
| MEK | 1 | ERK |
| SRC | 1 | p38 |
| MKK3/6 | 1 | p38 |
| MKK4 | 1 | p38 |
| RAS | 1 | JNK |
| FAK | 1 | ERK |
| PAK | 1 | MKK4 |
| MAP3Ks | 1 | MKK4 |
| EGFR | 1 | STAT1 |
| p38 | 1 | STAT1 |
| EGFR | 1 | STAT3 |
| SRC | 1 | STAT3 |
| EGFR | 1 | STAT5 |
| SRC | 1 | STAT5 |
| EGFR | 1 | SRC |
| PI3K | 1 | SRC |
| SERUM | 1 | MET |
| SERUM | 1 | PAK |
| SERUM | 1 | INSR |
| SERUM | 1 | EGFR |

**Supplementary Table 7.**

| PREDICTOR | TREATMENT | TIME [MIN] | FEATURE | DRUG | EFFECT SIZE |
| --- | --- | --- | --- | --- | --- |
| Median | EGF | 0 | Cas3<sup>cleaved</sup> | Pictilisib | -0.02739 |
| Median | EGF | 0 | GAPDH | Lapatinib | -0.02343 |
| Median | EGF | 0 | p-MKK4 | Pictilisib | 0.04569 |
| Median | EGF | 0 | p-p53 | Pictilisib | -0.02563 |
| Median | iMEK | 60 | p-AKT<sup>S473</sup> | Pictilisib | -0.04427 |
| Median | iMEK | 60 | p-ERK | Pictilisib | 0.042648 |
| Median | iMEK | 60 | p-MKK4 | Pictilisib | 0.05117 |
| Median | iMEK | 60 | p-p90RSK | Pictilisib | -0.0549 |
| Median | iPI3K | 60 | CyclinB | Lapatinib | 0.050149 |
| Median | iPI3K | 60 | IdU | Lapatinib | -0.03789 |
| Median | iPI3K | 60 | p-AKT<sup>T308</sup> | Lapatinib | 0.047966 |
| Median | iPI3K | 60 | p-MK2 | Lapatinib | -0.04315 |
| Median | iPI3K | 60 | p-MKK3/6 | Lapatinib | 0.040139 |
| Median | iPI3K | 60 | p-MKK4 | Fulvestrant | 0.041781 |
| Median | iPI3K | 60 | p-p90RSK | Pictilisib | -0.03437 |
| Cellular variability | EGF | 7 | p-S6 | Lapatinib | 0.02795 |
| Cellular variability | EGF | 7 | p-STAT3 | Lapatinib | -0.04125 |
| Cellular variability | EGF | 7 | p-STAT5 | Lapatinib | -0.06626 |
| Cellular variability | iPI3K | 40 | CyclinB | Fulvestrant | 0.032662 |
| Cellular variability | iPI3K | 40 | Ki-67 | Lapatinib | -0.05477 |
| Cellular variability | iPI3K | 40 | p-SMAD | Pictilisib | -0.04728 |
| Cellular variability | iPI3K | 40 | p-SRC | Pictilisib | 0.044885 |
| Cellular variability | iPI3K | 7 | p-AKT<sup>T308</sup> | Fulvestrant | 0.024168 |
| Cellular variability | iPI3K | 7 | p-MKK4 | Lapatinib | 0.022058 |
| Cellular variability | iPI3K | 7 | p-STAT3 | Pictilisib | -0.03228 |
| Edge flux | EGF | 0 | AKT<sup>T308</sup>.AKT | Lapatinib | -0.07514 |
| Edge flux | EGF | 0 | ERK.MK2 | Lapatinib | -0.06536 |
| Edge flux | EGF | 0 | MEK<sup>S221</sup>.MEK | Lapatinib | 0.048507 |
| Edge flux | EGF | 0 | mTOR.S6K | Lapatinib | -0.05844 |
| Edge flux | EGF | 0 | p38.MK2 | Lapatinib | 0.095079 |
| Edge flux | EGF | 0 | p90RSK.S6 | Lapatinib | -0.05934 |
| Edge flux | EGF | 0 | PI3K.BTK | Lapatinib | 0.052703 |
| Edge flux | EGF | 0 | SRC.BTK | Lapatinib | 0.059315 |
| Edge flux | EGF | 0 | AKT,RAS.MEK<sup>S221</sup> | Lapatinib | -0.05518 |
| Edge flux | iMEK | 60 | EGFR. STAT3 | Lapatinib | -0.06738 |
| Edge flux | iMEK | 60 | ERK.MK2 | Pictilisib | -0.08655 |
| Edge flux | iMEK | 60 | MKK3/6.p38 | Pictilisib | -0.04324 |
| Edge flux | iMEK | 60 | mTOR.S6K | Pictilisib | -0.04121 |
| Edge flux | iMEK | 60 | p38.p53 | Pictilisib | 0.06821 |
| Edge flux | iMEK | 60 | PI3K.BTK | Lapatinib | 0.047195 |
| Edge flux | imTOR | 7 | AKT<sup>T308</sup>.AKT | Lapatinib | -0.04717 |
| Edge flux | imTOR | 7 | MSK.NF-κB | Fulvestrant | 0.04297 |
| Edge flux | imTOR | 7 | PAK.MKK4 | Fulvestrant | 0.042304 |
| Edge flux | imTOR | 7 | PIP3.PDPK1 | Fulvestrant | -0.05599 |
| Edge flux | imTOR | 7 | PKC.GSK3β | Fulvestrant | -0.05353 |
| Edge flux | imTOR | 7 | SRC. STAT3 | Fulvestrant | -0.04018 |

